# Take ACTION to characterize the functional identity of single cells

**DOI:** 10.1101/081273

**Authors:** Shahin Mohammadi, Vikram Ravindra, David F. Gleich, Ananth Grama

**Author notes:** **Correspondence** All correspondences should be addressed to S.M. or A.G.

## Abstract

Single-cell transcriptomic data has the potential to radically redefine our view of cell type identity. Cells that were previously believed to be homogeneous are now clearly distinguishable in terms of their expression phenotype. Methods for automatically characterizing the functional identity of cells, and their associated properties, can be used to uncover processes involved in lineage differentiation as well as sub-typing cancer cells. They can also be used to suggest personalized therapies based on molecular signatures associated with pathology. We develop a new method, called *ACTION*, to infer the functional identity of cells from their transcriptional profile, classify them based on their dominant function, and reconstruct regulatory networks that are responsible for mediating their identity. Using *ACTION*, we identify novel Melanoma sub-types with differential survival rates and therapeutic responses, for which we provide biomarkers along with their underlying regulatory networks.

---

Complex tissues typically consist of heterogeneous populations of interacting cells that are specialized to perform different functions. A cell’s *functional identity* is a quantitative measure of its specialization in performing a set of *primary functions*. The *functional space* of cells is then defined as space spanned by these primary functions, and equivalently, the functional identity is a coordinate in this space. Recent advances in single cell technologies have greatly expanded our view of the functional identity of cells. Cells that were previously believed to constitute a homogeneous group are now recognized as an ecosystem of cell types [1]. Within the tumor microenvironment, for example, the exact composition of these cells, as well as their molecular makeup, have a significant impact on diagnosis, prognosis, and treatment of cancer patients [2].

The functional identity of each cell is closely associated with its underlying type [3]. A number of methods have been proposed to directly identify cell types from the transcriptional profiles of single cells [4–9]. The majority of these methods rely on classical measures of distance between transcriptional profiles to establish cell types and their relationships. However, these measures fail to capture weakly expressed but highly cell type-specific genes [10]. They often require user-specified parameters, such as the underlying number of cell types, which critically determine their performance. Finally, once the identity of a cell has been established using these methods, it is often unclear what distinguishes one cell type from others in terms of the associated functions.

To address these issues, we propose a new method, called *Archetypal-analysis for cell type identificaTION (ACTION)*, for identifying cell types, establishing their functional identity, and uncovering underlying regulatory factors from single-cell expression datasets. A key element of *ACTION* is a biologically inspired metric to capture cell similarities. The idea behind our approach is that the transcriptional profile of a cell is dominated by universally expressed genes, whereas its functional identity is determined by a set of weak but preferentially expressed genes. We use this metric to find a set of candidate cells to represent characteristic sets of *primary functions*, which are associated with specialized cells. For the rest of the cells, that perform multiple tasks, they face an evolutionary trade-off– they cannot be optimal in all those tasks, but they attain varying degrees of efficiency [11]. We implement this concept by representing the functional identity of cells as a convex combination of the primary functions. Finally, we develop a statistical framework to identify key marker genes for each cell type, as well as transcription factors that are responsible for mediating the observed expression of these markers. We use these regulatory elements to construct cell type-specific transcriptional regulatory networks.

We show that the *ACTION* metric effectively represents known functional relationships between cells. Furthermore, using the dominant primary function of each cell to estimate its putative cell type, *ACTION* outperforms state-of-the-art methods for identifying cell types. Furthermore, we report on a case study of cells collected from the tumor microenvironment of 19 melanoma patients [12]. We identify two novel, phenotypically distinct subclasses of *MITF*-high patients, for which we construct the transcriptional regulatory networks and identify regulatory factors that mediate their function. These factors provide novel biomarkers, as well as potential therapeutic targets for future development.

## Results

The *ACTION* framework consists of three major components, shown in Figure 1: (i) a robust, yet sensitive measure of cell-to-cell similarity, (ii) a geometric approach for identification of primary functions, and (iii) a statistical framework for constructing cell-type specific transcriptional regulatory networks (TRNs). Our framework starts by defining a cell similarity metric that simultaneously suppresses the shared but highly expressed genes and enhances the signal contributed by preferentially-expressed markers. The next component of our method is a geometric approach for identifying primary functions of cells. Each of these primary functions is represented by a corner of the convex hull of points defined within the functional space of cells. We refer to these corners as *archetypes* and the functional identity of each cell is represented by a convex combination of these archetypes. Finally, *ACTION* uses a novel method to orthogonalize archetypes, find key marker genes, and assess the significance of each transcriptional factor in mediating the transcriptional phenotype associated with each archetype. Finally, we use this method to construct the characteristic transcriptional regulatory network (TRN) of each cell type. In what follows, we describe, validate, and discuss each component in detail.

**Figure 1:**
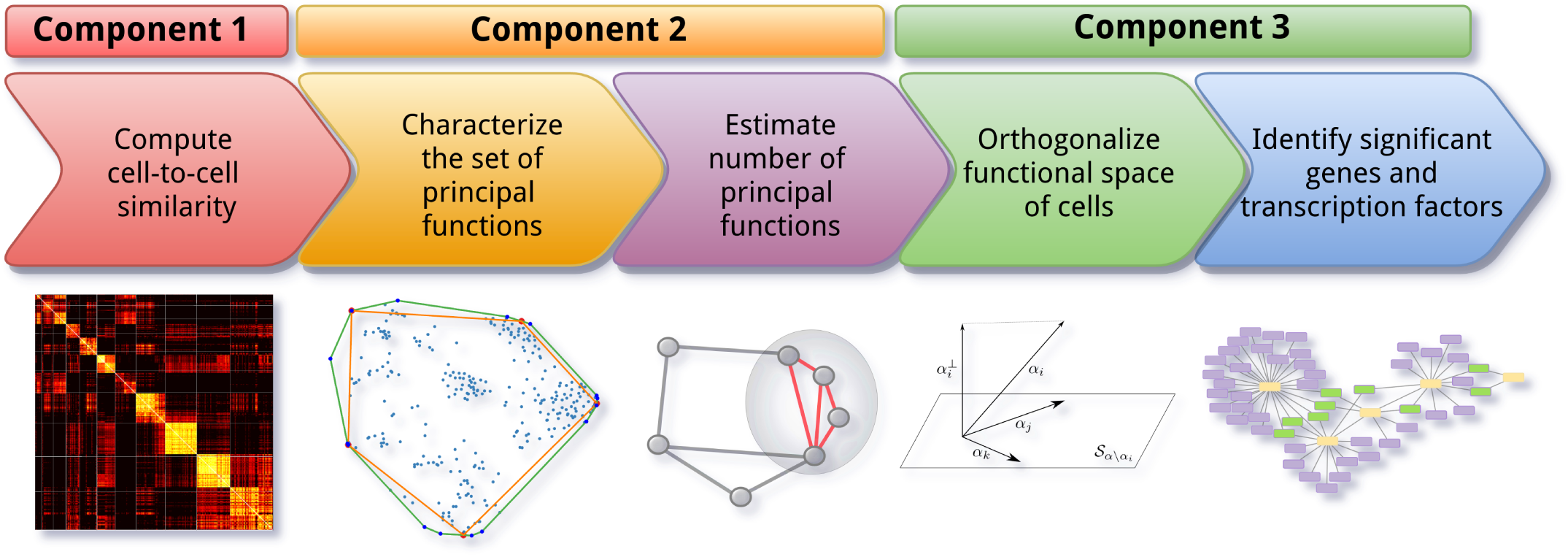
Overview of *ACTION*. *ACTION* consists of five main steps: (**i**) A biologically-inspired metric to capture similarity among cells. (**ii**) A geometric approach for identifying the set of primary functions. An automated mechanism for identifying the number of primary functions needed to represent all cells. An orthogonalization procedure for identifying key markers for each primary function. (**v**) A statistical approach for identifying key regulatory elements in the transcriptional regulatory network. These steps are grouped into three main components in the *ACTION* method that are each discussed in the methods section.

## The *ACTION* metric outperforms other metrics in representing functional relationships between single cells

A fundamental component of many methods for identifying cell types is a measure for quantifying the similarity between individual cells. Most prior methods rely on traditional measures, such as linear correlation that are not specifically targeted towards transcriptomic profiles. In contrast, we define a similarity metric, or formally a kernel, specifically designed for measuring the similarity between single cell transcriptomes [10]. Our approach is illustrated in Figure 2 and the mathematical models underlying the metric are described in the Methods section, Component 1. In summary, we first adjust the raw transcriptional profiles of cells to remove the effect of universally-expressed genes by projecting them onto the orthogonal space relative to the universally-expressed profile. We then boost the contribution of cell type-specific genes using an information theoretic approach. The final similarity is then a weighted inner-product kernel between these adjusted profiles.

**Figure 2:**
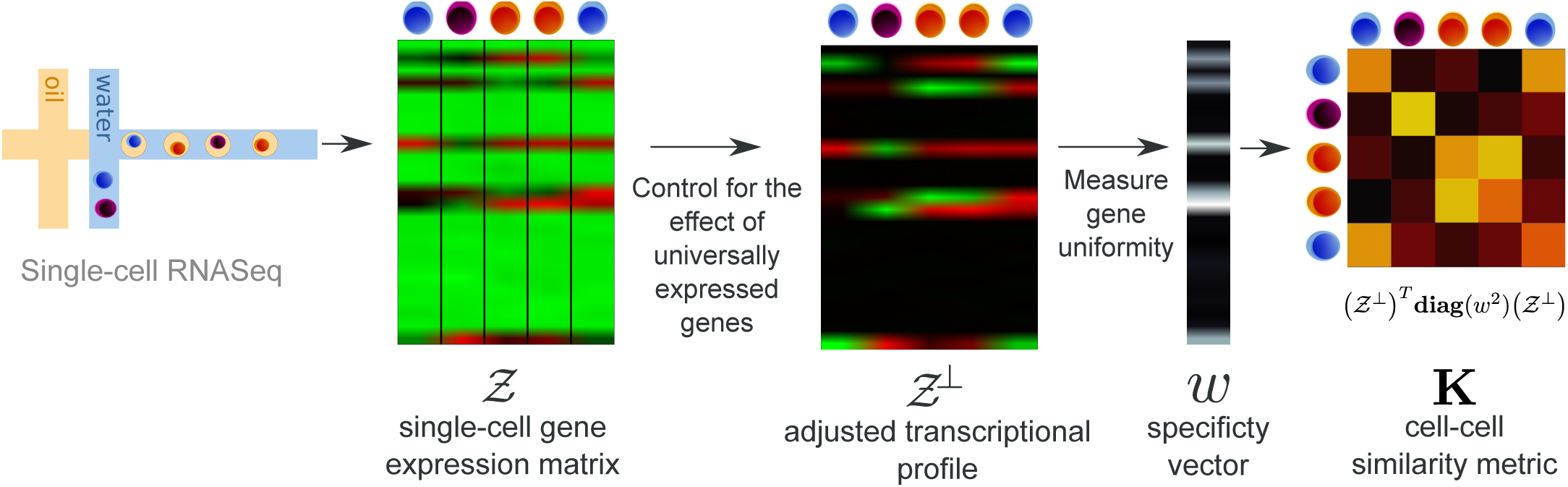
Workflow of *ACTION* cell-to-cell similarity metric. The *ACTION* metric is defined as a combination of two factors: (i) an adjusted transcriptional profile, in which the effect of universally-expressed genes has been masked out, and (ii) a gene specificity vector that assigns weights to each gene based on its informativeness. Finally, *ACTION* kernel is computed as the weighted dot-product of adjusted transcriptional vectors.

To establish the superiority of our metric, we compare it against an alternate measure specifically designed for single cell analysis, *SIMLR* [13]. SIMLR combines a number of distance metrics to learn a joint similarity score that maximizes the block diagonal structure of the resulting matrix. We also compare *ACTION* with the normalized dot product resulting from two nonlinear dimension-reduction techniques: *multidimensional scaling (MDS)* and *Isomap*. While *ACTION* is a non-parametric method, the other methods have one or more parameters that need to be specified by the user. For SIMLR, we need to specify the true number of cell types. For all methods other than *ACTION*, we must specify the dimension of the low-dimensional subspace. To give them the best chance at competing with *ACTION*, we evaluate ten different values for the dimension of projected subspace (from 5 to 50 with increments of 5) and report the best results obtained over all configurations.

To assess the quality of computed similarities between cells, we used each metric with kernel *k*-means, starting from 100 different initializations, in order to comprehensively assess their ability to identify discrete cell types. We apply this technique to four different datasets (see Methods, Datasets). These datasets are derived from different single cell technologies, have hundreds to thousands of cells, and span a wide range of normal and cancerous cells. We compare the predicted cell types against the annotated cell types in the original dataset using three different measures, namely *Adjusted Rand Index, ARI, F-score*, and *Normalized Mutual Information, NMI*.

Figures 3 present the performance of the cell type identification technique when operating with different similarity measures. In summary, our results demonstrate that in all cases the *ACTION* metric either outperforms or is jointly the best among competing metrics, except in the **Brain** dataset in which case *SIMLR* performs better when looking at all measures combined. A detailed analysis of the underlying distributions and the significance of differences among the top-ranked versus the runner-up methods is provided in the Supplementary Text 3. Additionally, for the **CellLines** dataset, which is specifically designed to evaluate cell type identification methods, we report the heatmap of marker genes for identified cell types to facilitate the visual assessment of the clustering differences, which is also available in Supplementary Text 4.

**Figure 3:**
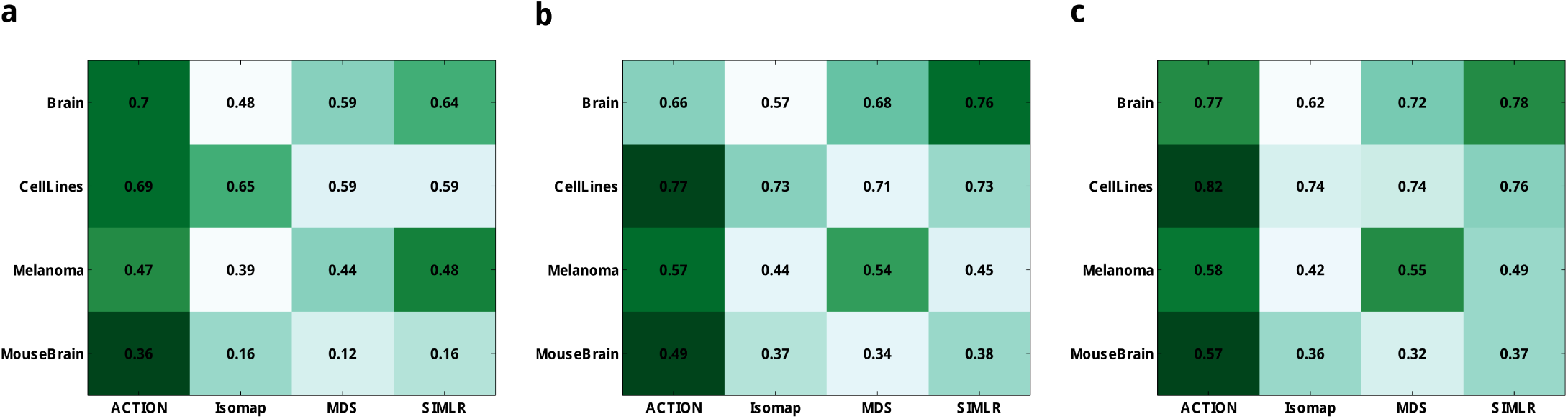
Performance of *ACTION* Similarity Metric. Various extrinsic measures of clustering quality for different cell similarity scores. (**a**) Adjusted Rand Index (ARI), (**b**) F-score, and (**c**) Normalized Mutual Information (NMI). All of these measures are upper-bounded by one with larger values indicating better results. The results in the table are the mean value over 100 individual runs of kernel k-means clustering with different initializations. The *ACTION* metric has no tunable parameters. For the other methods, we tested a range of parameters and report the best results. For each dataset, the corresponding row has been color-coded such that the darker green indicates better performance. Except for the Brain dataset, *ACTION* is either the best, or jointly the best. For Brain, the SIMLR metric is slightly better in an aggregation over all three measures.

To assess whether *ACTION* kernel can extract weak cell-type specific signals with increasing levels of dropout, we focus on the **CellLines** dataset that is specifically assayed to evaluate different cell type identification methods. We created a series of simulated expression profiles, seeded on the **CellLines** dataset, to mimic different levels of dropout. We iteratively removed nonzero elements at random, with the probability of removal being inversely proportional to the expression value, following previous work [14]. More specifically, the probability of removing each element is 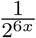, where *x* is the expression value. For each case, we generated 10 independent replicas and used each of them to compute different cell similarity metrics.Finally, we used each metric with kernel k-means and traced changes in the quality of clustering, which is presented in Figure 4. The *ACTION* method has the most stable behavior (RSS of the linear fit) with a minor downward trend as density goes below 10%. Furthermore, in each data point, *ACTION* has lower variation among different replicas. Other methods start to fluctuate unpredictably when density goes below 15%.

**Figure 4:**
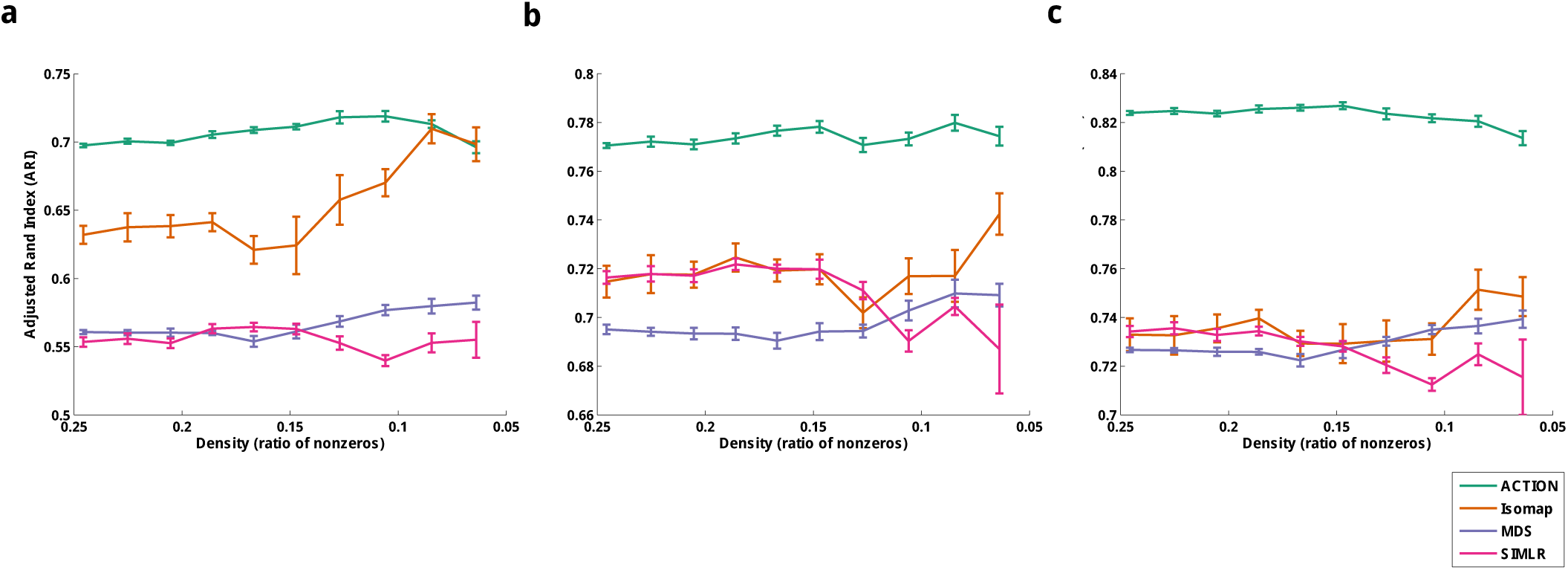
*ACTION* Kernel Robustness. A series of expression profiles with varying degrees of dropout has been simulated from the **CellLines** dataset. For each simulated network, we compute different metrics and use kernel k-means to identify cell types. The quality of cell type identification is assessed with respect to known annotation from the original paper using three different extrinsic measures. These results show that ACTION and MDS have the most stable performance over dropout.

**Figure 5:**
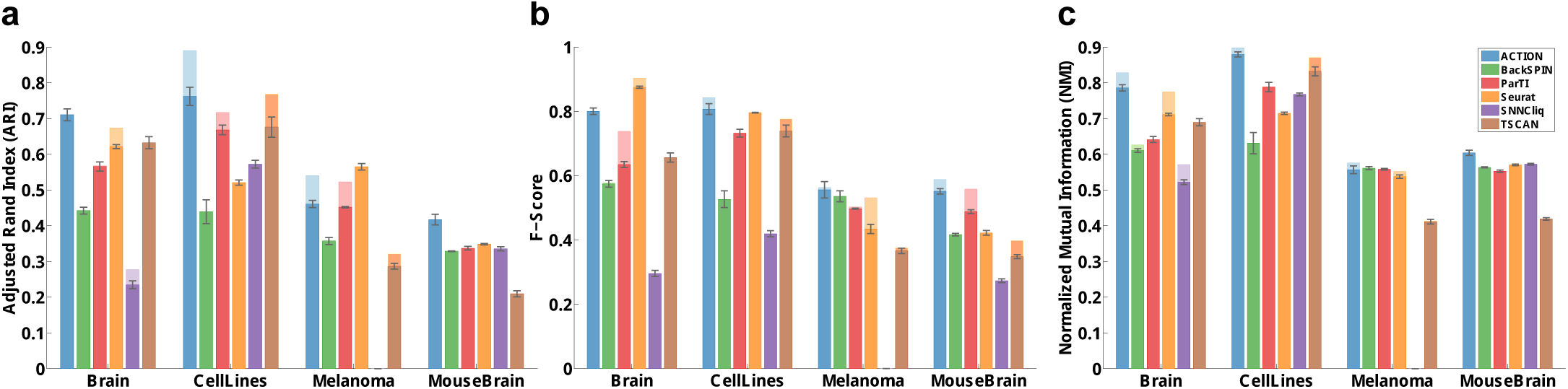
Performance of *ACTION* in identifying discrete cell types. *ACTION* identifies cell types by classifying cells according to their dominant primary function (closest archetype). Performance is measured via various measures with respect to the cell types provided with the data: (**a**) Adjusted Rand Index (ARI),(**b**) F-score, and (**c**) Normalized Mutual Information (NMI) of cell type identification. Larger values are better, and the perfect score (upper bound) is one. Lighter shades are the actual results when using all cells/ samples, whereas the darker bar and the error bar indicates a 10-fold test to estimate the variability and stability of predictions for each method. In the **CellLines** dataset, which was originally created to benchmark cell type identification methods, *ACTION* outperforms other methods with respect to ARI and NMI measures, and ties with *Seurat* in terms of F-score. In the **MouseBrain** dataset, *ACTION* significantly outperforms other methods in all three measures. In the **Brain** datasets there is a competition between *ACTION* and *Seurat*, whereas in the **Melanoma** there is more variability among different methods. This is particularly associated with the level of annotations in this dataset (lack of annotations for T-cell subclasses and tumor subtypes, for example) and the varying resolution of different methods.

Overall, these results establish the *ACTION* metric as a *fast, nonparametric*, and *accurate* method for computing similarity among single cells. We use this measure throughout the rest of our study.

## The *ACTION* method successfully uncovers functional identity of single cells

Using the *ACTION* metric as a measure of similarity between cells, we develop a new method for characterizing the *functional identity* of cells in a given experiment. Our method is based on a geometric interpretation of cellular functions. In this view, each cell corresponds to a data-point in a high-dimensional space. Our method identifies *“extreme”* corners, or *archetypes* in this space, each of which represents a *primary function*. The functional identity of each cell is subsequently characterized as a convex combination of these primary functions. (A convex combination is a linear combination of points, such that all coefficients are non-negative and sum to 1.) The choice of the number of primary functions or archetypes is based on a novel non-parametric statistical procedure. See Methods section, Component 2 for a detailed description.

### Discrete view of cell types

To approximate discrete cell types from the primary functions identified using *ACTION*, we assigned each cell to a single dominant function, as determined by its closest archetype. We compare our method to five recently proposed methods: Seurat (v2.2) [15], SNNCliq [7], BackSPIN [16], single-cell ParTI [8, 17], and TSCAN [9] (see Supplementary Text 1 for a brief description of these methods) to predict annotated cell types on the same four datasets (see Methods, Datasets). For the *Melanoma* dataset, *SNNCliq* did not terminate after *72* hours, after which we stopped the experiment.

We report the results of each method applied to each dataset. In addition, to further validate these results, we select 90% of cells in each dataset, proportional to the total cell type counts, and run each method on each of these 10-folds, and report mean and standard deviation of these results. In all cases, we observe that *ACTION* performs as well or better than the other methods. For the **Melanoma** dataset, however, there is no consensus among the top-ranked methods. This can be attributed, in part, to the extent of available annotations in this dataset and the varying resolution of different methods. We further investigate our results on this dataset in the following sections.

In terms of computational time, graph-based techniques, such as *SNNCliq* and *Seurat*, perform better than *ACTION* for smaller datasets; however, *ACTION* scales more gracefully as the size of the dataset increases (see Supplementary Text 8 for the details). Also, an example heatmap for the results of the **CellLines** dataset is provided in the Supplementary Text 5 for an illustration of the benefits of our approach.

In Supplementary Text 9, we study the robustness of *ACTION* in presence of noise and outliers, as well as its sensitivity to identify rare cell types. We found that preconditioning the adjusted expression profiles significantly enhances the accuracy of predictions, while relaxing the pure pixel assumption further stabilizes these predictions. Furthermore, we show that our method is sensitive enough to identify rare cell types with 2% of the total population. Below this population, they are characterized as noise and outlier cells.

Overall, these experiments show that *ACTION*, while designed to explore the continuous functional space of cells, is successful in identifying discrete cell types as stable states in this space.

### Continuous view of cell states

While the functional identities of cells can be discretized to define cell types, they can also be explored in the continuous space of all primary functions. To illustrate this *continuous view*, we perform a case study on the *Melanoma* dataset (Figure 6). Each point corresponds to a cell. Given the functional profile of cells, defined in a *k*-dimensional space, with *k* being the number of archetypes, we map cells to a two-dimensional plane using the Stochastic Neighbor Embedding (SNE) method with a deterministic initialization (see Supplemental Text 10). Our non-parametric method selected 8 archetypes for the Melanoma data, each is marked with a text label (A1,…, A8) and assigned a unique color. We interpolate the color of each cell using its distance from all archetypes to highlight the continuous nature of the data. We use markers from *LM22* dataset [18] to distinguish different subtypes of T-cells. For the tumor cells, we perform further analysis of active transcription factors, as described in the next section and the methods section, to identify key driving regulators that distinguish each archetype.

**Figure 6:**
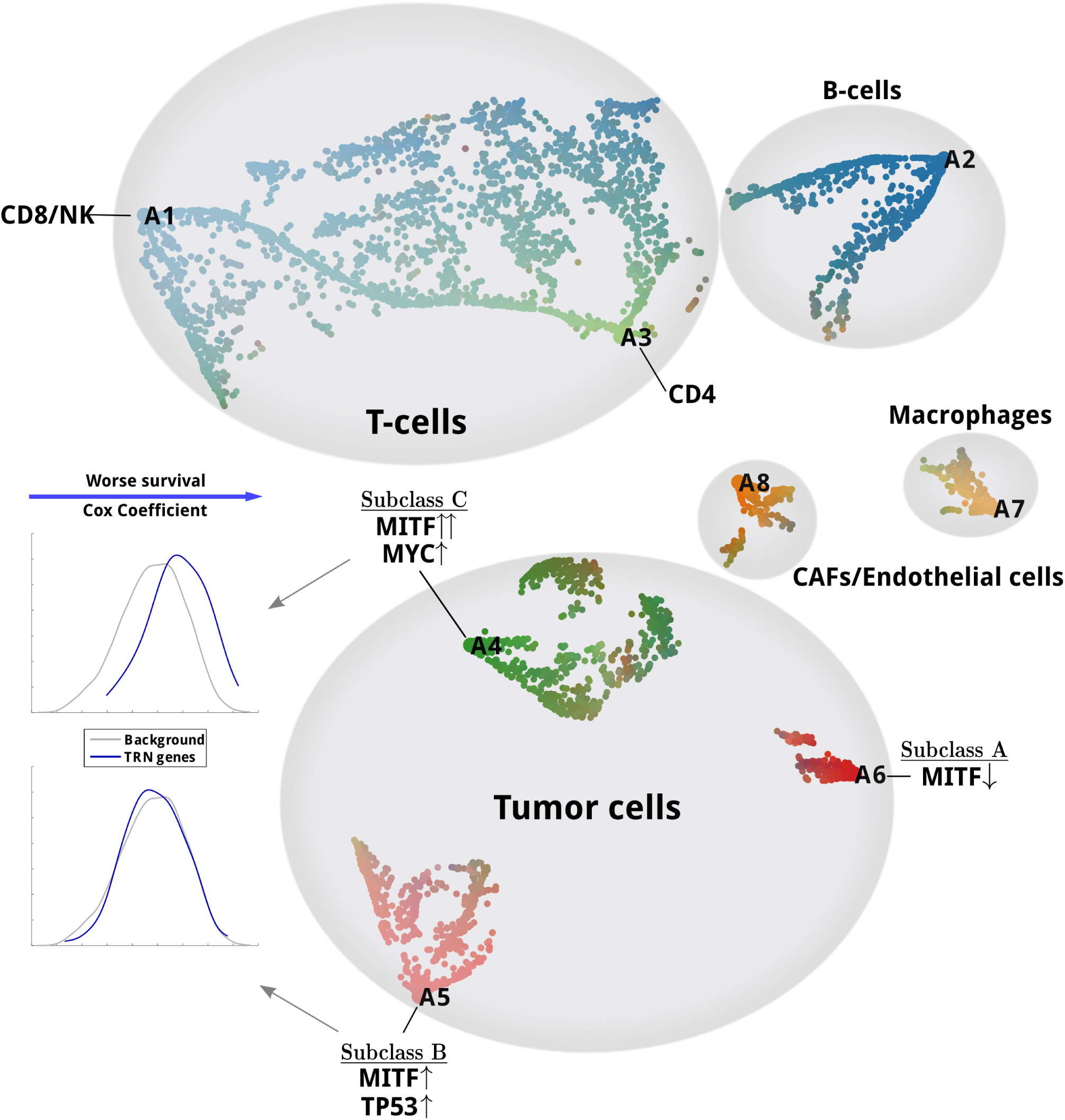
A continuous view on the space of primary functions in the Melanoma dataset. Each archetype, representing a primary function, is illustrated using a textual label (A1-A8). Each small dot represents a cell. Cells are color coded based on their proximity to archetypes. All data points are projected onto a 2D plane using a carefully initialized Stochastic Neighbor Embedding method (SNE, see Supplemental Text 10). The functional space of cells exhibit a mix of cell state continuum, such as in the case of T-cells, as well as discrete cell types. Three subclasses of melanoma tumor cells are marked accordingly in the map. Subclasses *B* and *C* are both MITF-associated. Among them, genes that participate in the transcriptional regulatory network (TRN) for *subclass B* do not show any significant shift in Cox coefficient, compared to the background of all genes, whereas in *subclass C* they do. In this sense, high-expression of genes in the TRN of *subclass C* is significantly associated with worse outcome in the melanoma patients.

Figure 6 demonstrates the ability of our method to identify both isolated cell-types with specialized primary functions, as well as the ones with a mixed combination of functions. As an example, T-cells constitute a continuous spectrum across functional space of cells, which is consistent with previous studies [19].Subclasses of melanoma cells, on the other hand, exhibit distinct separation and have unique phenotypic behaviors and survival rates. In what follows, we identify key marker genes for each subclass, transcription factors that are significantly associated with regulating these genes, and construct their gene regulatory network.

## The *ACTION* method constructs accurate models of the regulatory networks that drive functional identity of cells

We propose a new method for constructing regulatory pathways responsible for mediating the phenotypes associated with each archetype. We first perform an *archetype orthogonalization* to compute a residual expression and identify marker genes that are unique to each archetype. We then assess the role of each transcription factor (TF) in controlling these marker genes. Significant TFs, together with their top-ranked target genes (TGs), constitute the underlying transcriptional regulatory network (TRN) that is responsible for mediating a given primary function, and consequently, the phenotype associated with cells dominantly associated with that function (see Methods, Component 3, and Figure 7a for additional details).

**Figure 7:**
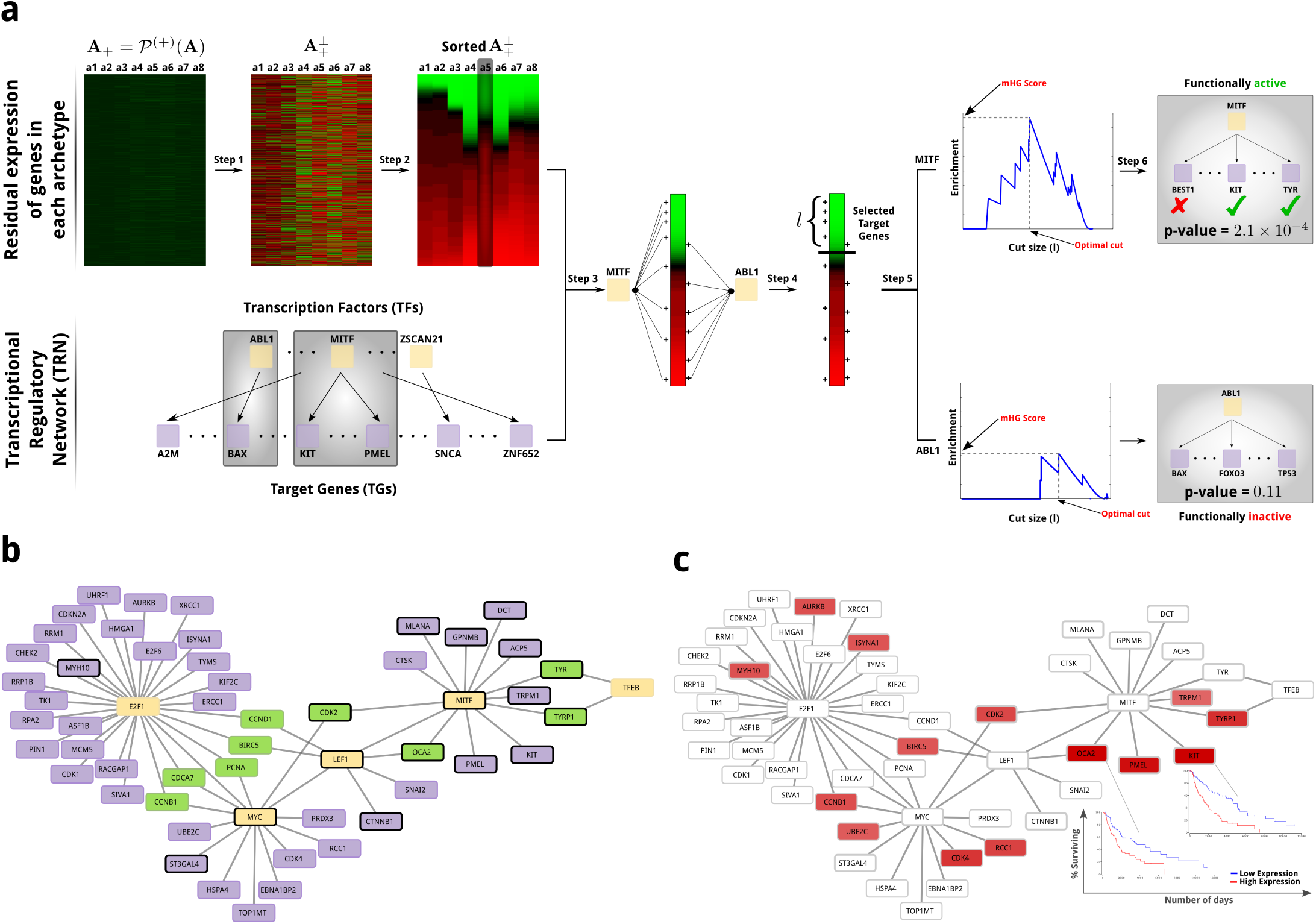
The transcriptional regulatory network (TRN) for MITF-associated Melanoma patients highlights a number of genes that have not previously been associated with Melanoma – along with some known markers. (**a**) Main steps involved in the construction of archetype-specific TRNs: (1) Orthogonalize archetypes with respect to each other, (2) Sort genes based on their residual expression, (3) Map gene targets for TFs to the sorted list genes, (4) Enrichment analysis for fixed cut size *l*, (5) Find optimal cut size and compute minimum HyperGeometric (mHG) score, and (6) Assess significance of the mHG score using Dynamic Programming (DP). (**b**) A subset of the TRN of *subclass A* induced by using only the most significant TFs. The yellow nodes are transcription factors (TF), the purple nodes are target genes (TG), and green nodes are target genes that bridge different TFs. Genes marked with black border are known to be involved in the proliferative subclass of Melanoma. (**c**) The TRN of *subclass A* with genes color-coded according to their Cox coefficient. Red genes are the ones whose high expression is associated with worse outcome, and brightness of the color relates to the severity of the outcome. Kaplan-Meier plots for two of the targets of MITF that are unique to *subclass A* but not *subclass C* are shown on the plot.

To evaluate the quality of top-ranked genes identified after orthogonalizing each archetype, we selected the top 20 genes and marked the ones that are known markers (according to the original paper) for the cell type that is enriched for the archetype. Supplementary Text 11 presents a complete table of these top-ranked genes, where known marker genes are in bold typeface. Upon initial observation, a large fraction of these genes appear to be associated with known markers. To systematically assess the significance of this event, we created a label vector for each archetype according to its sorted list of genes after orthogonalization. Then, we use mHG *p*-value to assess the enrichment of markers among top-ranked genes, which are presented in the last row in the table. It is notable that all archetypes are highly significant with respect to the enrichment of marker genes among their top-ranked residual genes, with the exception of CD4 T-cell and tumor subclass A. After further examination, we observed that the majority of T-cell markers provided in the paper are for CD8 T-cell and provided tumor markers in this dataset are for MITF over-expressed melanoma tumors. Thus, the corresponding columns have less significant results that the others.

Next, to distinguish different subclasses of tumor cells, we computed the transcription factors that are significantly associated with regulating the top-ranked marker genes for each archetype, as well as the particular subset of target genes that they regulate. We found that both subclasses *B* and *C* are associated with *SOX10* and *MITF*, two of the most well-characterized markers for “proliferative” melanoma tumors [20]. Further analysis of these factors, however, reveals that while both of these subclasses are MITF-associated, the degree of association is higher for *subclass C*. Examining downstream targets of MITF that are activated in each subclass (see Supplementary Text 13), we identified that *GPNMB, MLANA, PMEL*, and *TYR* are shared between two subclasses, whereas *ACP5, CDK2, CTSK, DCT, KIT, OCA2* and TRPM1/P1 are unique to *subclass C*. To validate these targets, we used a comprehensive list of down-regulated genes in response to MITF knockdown in 501Mel melanoma cells [21]. The overlap of identified MITF target genes and the set of down-regulated targets was significant for subclasses *B* and *C* (hypergeometric test *p*-values of 7.5 *×* 10^−5^ and 1 *×* 10^−6^, respectively). This further validates that our method is identifying not only the right transcription factors, but also the right set of target genes for them. Among other distinguishing TFs, subclass *B* is significantly associated with *BRCA1* and *TP53*, whereas subclass *C* is associated with *MYC*. Factors *BRCA1* and *TP53* are both tumor-suppressors, whereas *MYC* is a proto-oncogene. Activation of these transcriptional factors, in turn, can differentially regulate downstream targets that may contribute to worse outcome in subclass *C*.

Based on these observations, we propose the hypothesis that subclass *C* should have worse outcome than subclass *B*. To support this hypothesis, we construct subclass-specific transcriptional regulatory networks (TRN) for these two subclasses. (The complete TRNs for each of the 8 archetypes are available for download, see Supplemental Text 14). The set of transcription factors in these networks have a total of 51 and 91 distinct target genes, respectively, that are functionally active. In order to understand how the difference among these genes contribute to the overall survival of patients, we assessed the association between identified genes in each network and survival rate of Melanoma patients in the TCGA dataset, measured via multivariate Cox regressions [22]. We note that genes in *subclass C* significantly deviate from the null distribution of Cox coefficients for all genes (Kolmogorov-Smirnov test; *p*-val = 5.4 *×* 10^−10^), whereas genes in *subclass B* do not (*p*-value = 0.31), which translates into worse prognosis for *subclass C*. These observations are summarized in Figure 6.

To further study the underlying regulatory mechanisms that drive this poor-outcome phenotype for subclass *C*, we focus on only the most significant transcription factors (those with functional activity *p*-values ≤ 10^−3^, rather than ≤ 0.05 above) and construct their associated regulatory network. Figure 7a shows the interaction network among highly significant TFs and their major targets in *subclass C*. While some of these factors, and their target genes, were previously directly or implicitly associated with Melanoma, this network provides a novel systems view of the interactions, and highlights new regulatory interactions. For instance, amplification of the *MYC* oncogene has been long associated with poor outcome in Melanoma patients [23]. Also, *E2F1* is a critical transcription factor that is involved in cell cycle transition from G1 to S phase, and its overexpression is commonly associated with poor patient survival in high-grade tumors [24]. The *LEF1* factor has a dual role. On one hand, it acts as a downstream effector of the Wnt signaling pathway and is associated with phenotype switching in Melanoma cells between proliferative and invasive states [25]. On the other hand, it has been suggested that *LEF1* has a distinct, Wnt-independent, role in activating *E2F1* [26]. Finally, we note that *LEF1* regulates both *MITF* and *MYC*. Collectively, we hypothesize that *LEF1* is a key TF that regulates phenotype switching from proliferative to invasive state in *subclass C*, by controlling other transcription factors, including *MITF, MYC*, and *E2F1*.

To revisit the problem of survival analysis, and to recover genes that affect this prognostic change, we project individual Cox coefficients for each gene onto the TRN of *subclass C* (Figure 7b). Two of the most significantly associated genes, *KIT* and *OCA2*, are among *MITF* targets that are unique to *subclass C* but not *subclass B*. The Kaplan-Meier plots for these two genes are visualized alongside the TRN. In addition, there are multiple targets of *MYC, LEF1*, and *E2F1* that are also associated with poor outcomes for melanoma patients.

Finally, to assess the therapeutic indications of these subclasses, we used the pharmacogenomic profiling of cancer cell lines [27]. There are 53 melanoma cell lines in this dataset. For each of these cell lines, we have access to both their transcriptomic profile and drug response for 256 different drugs. We used the top 100 genes in subclasses *A-C* to find cell lines that closely resemble each of these subclasses. We *z*-score normalize each row of this submatrix and use mHG *p*-value to assess the the enrichment of marker genes among top ranked genes. We use a *p*-value of 10^−3^ to ensure that selected cell lines are closely related to original subclasses. This leaves us with 9, 6, and 15 cell lines for subclasses *A, B* and *C*, respectively, and 23 unclassified cell lines. For cell lines associated with subclasses *B* and *C*, we used a *t*-test to assess differences in the distribution of IC50 value between these two subclasses. We found that subclass *C* is more sensitive to the drugs targeting ERK MAPK signaling, specifically *Refametinib, CI-1040, PLX-4720, SB590885, Selumetinib, AZD6482, PLX-4720*, and *Dabrafenib*, among which *PLX-4720* and *Dabrafenib* are the most effective ones.

## Methods

### Datasets

#### Single cell gene expression datasets

For all our studies, we rely on the following datasets collected from publicly available sources:

#### *Brain* (GEO: GSE67835)

This dataset contains 466 cells spanning various cell types in the human brain, including astrocytes, oligodendrocytes, oligodendrocyte precursor cells (OPCs), neurons, microglia, and vascular cells [28].

#### *CellLines* (GEO: GSE81861)

This dataset is recently published to benchmark existing cell type identification methods. It contains 561 cells from seven different cell lines, including A549 (lung epithelial), GM12878 (B-lymphocyte), H1 (embryonic stem cell), H1437 (lung), HCT116 (colon), IMR90 (lung fibroblast), and K562 (lymphoblast). To assess the effect of batch effects, GM12878 and H1 are assayed in two batches [29].

#### *Melanoma* (GEO: GSE72056)

This dataset measures the expression profile of 4,645 malignant, immune, and stromal cells isolated from 19 freshly procured human melanoma tumors. These cells are classified into 7 major types [12].

#### *MouseBrain* (GEO: GSE60361)

This dataset contains the expression profile of 3005 cells from the mouse cortex and hippocampus. These cells classify into seven major types, including *astrocytes-ependymal, endothelial-mural, interneurons, microglia, oligodendrocytes, pyramidal CA1*, and *pyramidal SS* [16].

#### Transcriptional Regulatory Network (TRN)

We collect transcription factor (TF) – target gene (TG) interactions from the TRRUST database [30]. This dataset contains a total of 6, 314 regulatory interactions between 651 TFs and 2, 102 TGs.

#### Drug sensitivity in cell lines

We downloaded processed gene expression and drug sensitivity data from the Genomics of Drug Sensitivity in Cancer Project website [27]. This datasets consists of a total of 1,001 cell lines, spanning different types of cancer, 52 of which are melanoma cell lines that also have their gene expression profile available. A total of 256 compounds were screened on these cell lines IC59 values for each pair has been reported.

### Component 1: Computing a biologically-inspired metric to represent functional relationships between cells

The transcriptome of each cell consists of genes that are expressed at different levels and have different specificity with respect to the underlying cell types. *Universally-expressed genes* correspond to the subset of genes responsible for mediating core cellular functions. These functions are needed by all cells to function properly, which result in ubiquitous expression of these genes across all cell types [31]. While fundamental to cellular function, these genes are not informative with respect to the identity of cells. On the other hand, cell type-specific genes are preferentially expressed in one or a few selected cell types to perform cell type-specific functions. Unlike universally-expressed genes, cell type-specific genes are, typically, weakly expressed, but are highly relevant for grouping cells according to their common functions. Our goal here is to define a similarity measure between cells that suppresses universally expressed genes and enhances the signal contained in cell type-specific genes.

#### Suppressing universal but highly expressed genes

To suppress the ubiquitously high expression of universally-expressed genes, we adopt a method that we developed recently for bulk tissue measurements and extend it to single cell analysis [10]. This method projects a standardized representation of expression profiles of cells onto the orthogonal subspace of universally-expressed genes. Let us denote the *raw expression profile* of cells using matrix **Χ** *∈* ℝ^*m×n*^, where each row corresponds to a gene and each column represents a cell. We use ***x***_*i*_ to denote the expression profile of *i*^*th*^ cell. In addition, let us denote the signature vector of universally-expressed genes by *υ*. As a first order estimate, a universally-expressed signature is computed by taking the average expression over all cells: 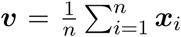; that is, *υ*_*i*_ is the average expression of gene *i* across all samples. This choice is motivated by the fact that highly expressed genes are more consistently expressed, whereas lowly expressed genes show exhibit higher variability. To this end, by orthogonalizing with respect to the mean value, we significantly reduce the effect of universally expressed genes, while preserving the variation of lowly-expressed, but preferential ones [32]. After estimating this baseline expression, we *z*-score normalize the profile of each cell: 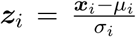, where *μ*_*i*_ and *σ*_*i*_ are the mean and sample standard deviation of the entries in the *i*th cell profile. Similarly, we *z*-score normalize the signature vector of universally-expressed genes, υ, to create a new vector *z*_*υ*_. Finally, we project out the impact of the universally-expressed gene expressions on each cell’s profile as follows:

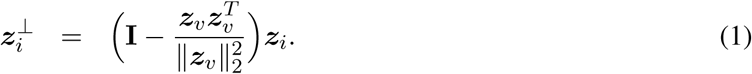

This operation projects *z*_*i*_ to the orthogonal complement of the space spanned by the universally-expressed genes. We then concatenate the column vectors 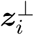 to create a *adjusted cell signature* matrix *Ƶ*^⊥^

#### Enhancing signal from cell type-specific genes

Next, to enhance the signal contributed by preferentially expressed genes, we propose an information theoretic approach that is inspired by the work of Schug *et al.* [33]. The main idea is to use Shannon’s entropy to measure the informativeness of genes. If a gene is uniformly utilized across cells, it contains less information as opposed to the case in which it is selectively active in a few cells. To this end, we start with the positive projection of adjusted cell signatures, 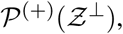 in which case we replace all negative values with zeros. Then, we normalize this matrix to construct a stochastic matrix **P** (every row sums to one). Let *p*_*i*_ be the row vector associated with the *i*^*th*^ gene. We compute the uniformity, or normalized entropy, of *p*_*i*_ as: u(i) = – ∑_*j*_ *p*_*ij*_ log(*p*_*ij*_)/log(*n*), where *p*_*ij*_ is an entry in the matrix **P** and *n* is the number of genes. This value is always between zero and one and is used as a basis to boost contributions from the most informative genes. A detailed comparison of our entropy-based method with dispersion and Gini index is provided in the Supplementary Text 2.

To scale genes according to their specificity, we compute a coefficient that controls the contribution of each gene. This coefficient is greater than one (scales up) for cell type-specific genes and less than one (scales down) for universally expressed genes, respectively. To do so, we note that the distribution of the entropy values follows a bimodal distribution, with separate peaks for the cell type-specific and universally-expressed genes. To identify the critical point where these two population separate from each other, we fit a mixture of two Gaussians over the distribution of the values and use it to identify this transition point, denoted by *u\**, which is the point of equal probability from each Gaussian.Then for each gene *i*, we define a scaling factor as *w*_*i*_ = *u*/u*(*i*). Finally, we compute the kernel matrix as follows:

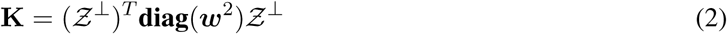

In this formulation, if we denote **Y = diag**(*w*)*Ƶ*^⊥^ then **K** is a dot-product kernel defined as **Y**^*T*^ **Y**. We will refer to **Y** as the *adjusted transcriptional profile* of cells, and **K** as the *cell similarity kernel*, or *ACTION metric*.

### Component 2: Fitting a geometric construct to characterize functional identity of cells

Due to evolutionary constraints, biological systems, including cells, that need to perform multiple *primary functions*, or tasks, can not be optimal in all those tasks; thus, these systems evolve to be specialized in specific tasks [11]. The functional space of cells then can be represented by a low-dimensional geometric construct, such as a polytope, the corners of which correspond to the set of specialized *primary functions*. The convex hull of a given set of points is the minimum volume polytope that encloses all points. This can be envisioned as a rubber band fitting to the outermost points. Constructing the convex hull in high-dimensional space is computationally expensive and susceptible to noise and overfitting. As an alternative, we seek a limited number of points on the convex hull that enclose as many points as possible, while being resilient to noise and outliers. Each point here represents a cell and each corner, or archetype, of this polytope is a candidate cell that best represents a unique *primary function*. To find these candidate cells, we use a modified version of the *successive projection algorithm (SPA)* combined with a novel model selection technique to identify an optimal number, *k*, of candidate cells on the approximate convex hull that best represent distinct pure cells with specialized primary functions. Finally, we use the *principal convex hull algorithm (PCHA)* to relax these corners to allow others cells to contribute to the identity of each archetype/corner.

#### Identifying the best *k* representative cells

Formally, given a matrix **Y** representing the *adjusted transcriptional profile* of cells, we aim to construct an optimal set *𝒮* of *k* columns such that each selected column is an ideal representative of the cells that perform a given primary function. Let us assume that matrix **Y** can be decomposed as **Y** = **Y**(:, *𝒮*)**H** + **N**, where *𝒮* is the selected column subspace of matrix **Y**, **H** is nonnegative with column-sums equal to one, and **N** represents bounded noise, where ||**N**(:,*j*)||_2_ ≤ *∊* That is, we can select |***𝒮***| = *k* columns from matrix **Y** to represent rest of the columns, with consideration for noise. A matrix satisfying this condition is called *near-separable* and is known as the *near-separable nonnegative matrix factorization (NMF)* when **Y** is nonnegative. For a matrix satisfying near-separability, there is an efficient algorithm, with provable performance guarantees, that can identify columns in *𝒮*. Furthermore, premultiplying matrix **Y** with a nonsingular matrix **Q** preserves its separability, but if chosen carefully, can enhance the conditioning of the problem and accuracy of results. To find the optimal preconditioning matrix **Q**, we use a theoretically-grounded method based on identifying a minimum volume ellipsoid at the origin that contains all columns of **Y** (Supplementary Text 6).

#### Estimating the optimal number of cells to represent primary functions

Given that *SPA* selects *k* columns of **Y**, *given k*, the next issue is how to find the optimal value of *k* that captures most variation in data without overfitting. We devised a novel monitoring technique that assesses the current *k*-polytope to see if there is any evidence of oversampling the cell-space. If so, it stops the algorithm. Otherwise, it continues by adding new archetypes. Informally, oversampling happens when we start adding new archetypes to regions in the space that are already well-covered by other archetypes, in which case the newly added archetype would be significantly close to one or more other archetypes, compared to the rest of the archetypes. Given that each archetype is a candidate cell, we can measure relationship between them using the *ACTION* metric. The distribution of similarities resembles a normal distribution; however, as we start to oversample, the right tail of the distribution starts getting heavier. To distinguish the pairs of archetypes in this heavy-tailed region, we *z*-score normalize pairwise similarities between archetypes and select all pairs whose *z*-transformed similarity scores are above 1.96, which corresponds to 95% confidence level under Gaussian assumption for the underlying distribution. Then, we build an *archetype similarity graph* using these pairs of close archetypes. In this graph, oversampling can be identified by the emergence of dense local regions. We use the Erdős-Rényi (ER) random graph model as a background to assess density of each sub-region, or connected component, in the archetype similarity graph [34, 35]. If we find at least one of the connected components that is significantly dense, which is a sign of oversampling, then we terminate the algorithm and choose the last value of *k* before oversampling happens.

#### Optimizing archetypes by relaxing the pure cell assumption

After estimating *k* ideal candidate cells, or pure cells, we use archetypal-analysis (AA) [36], which can be viewed as a generalization of near-separability to relax corners by locally adjusting them to have contributions from multiple cells. Formally, we can formulate *AA* as follows:

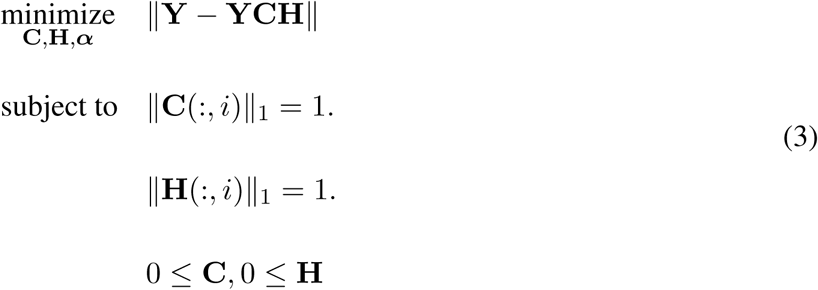

Near-separable non-negative matrix factorization is a special case of AA in which **Y** is non-negative, **C** has exactly *k* nonzeros, and none of the columns have more than one element. We use an efficient algorithm, called *Principal Convex Hull Analysis (PCHA)*, to solve the above problem to a local optima.

The matrix **A** = **YC** then stores the *archetypes*. Column stochasticity of **C** indicates that archetypes are convex combinations of data points, and column stochasticity of **H** indicates each data point can be represented as convex combination of archetypes.

A complete pseudo-code fitting all these components together is provided in Supplementary Text 7.

### Component 3: Constructing the driving transcriptional regulatory network for each archetype

In order to understand what control mechanisms are responsible for mediating the transcriptional phenotype of each archetype, we first have to identify key marker genes that distinguish a given archetype from the rest of archetypes (see Figure 7a for an illustrative guide to this section). To this end, we first orthogonalize each archetype with respect to all other archetypes. In this formulation, what remains, referred to as the *residual expression* of genes, ranks genes according to their importance in a given archetype. Let matrix **A** = **YC** represent the identified archetypes. Let **A**^(+)^ = *𝒫*^(+)^(**A**) be the projection to positive entries and let 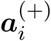 stand for the column *i* of **A**^(+)^. Moreover, let 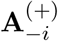 denote the matrix without the *i*th column. Our goal is to project 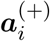 into the subspace orthogonal to the columns spanned by 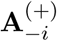. Then, the orthogonalization step can be written as:

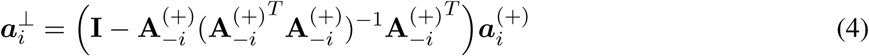

Finally, we construct matrix 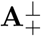 where each column is 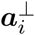. Terms in this matrix are called *residual expressions* and help identify distinguishing marker genes for each archetype.

Those genes with high residual expression in each archetype are controlled through regulatory networks within the cell. To uncover these relationships, we identify transcription factors that are significantly associated with the expression of marker genes, which we will refer to as *functionally active* TFs. Functional activity of TFs is inferred directly from the expression of their target genes; thus, these TF activities can be controlled at different stages, ranging from transcriptional to post-translation regulations. To infer these activities, we first need to classify their target genes as either active or inactive in a given context (archetype). We partition genes according to their residual expression and declare top-ranked genes as active. We use the minimum hypergeometric (mHG) method [37] to find the optimal partition of genes and assign a *p*-value to it. The main step of this algorithm is similar to classic enrichment analysis: for a fixed size *l*, we use the hypergeometric *p*-value to assess the over-representation of target genes for a given TF among top‐*l* markers for an archetype. Then, we compute the same statistic for all 1 ≤ *l* ≤ *m*, where *m* is the total number of genes. The minimum hypergeometric tail that is obtained, referred to as the *mHG score*, specifies the best cut, *l*^(*best*)^, and all target genes that are ranked higher than *l*^(*best*)^ among marker genes are selected as regulated targets for that TF. Finally, we use the obtained *mHG score* to assess the significance of the TF itself. This can be accomplished using a dynamic programming algorithm that assesses the probability of observing the same or more significant mHG score within the population of all binary vectors of size *m* with exactly *r* nonzeros, where *r* is the number of targets for the current TF. The set of all significant transcription factors (TFs), together with their target genes (TGs) that fall above the cut that results in the mHG score, are used to construct the final transcriptional regulatory network (TRN).

## Data Availability

All materials and codes are readily available for download from http://compbio.mit.edu/ACTION.

## Acknowledgements

This work is supported by the NSF Center for Science of Information STC (CCF-0939370), NSF Grants BIO 1124962, NSF IIS-1546488, NSF CCF-1149756, IIS-1422918, the DARPA SIMPLEX program, the Sloan Foundation, and NIH Grants 5R01AI114814-02 and 5U01CA198941-03.

## Author Contributions

SM conceived the idea, designed and implemented the method, ran the experiments, analyzed the results, and drafted the manuscript. VR performed initial experiments that led to the final method. DG and AG helped design the method, analyzed results, and assisted with the writing. All authors read and approved the final manuscript.

## Competing Interests

The authors declare that they have no competing financial interests.

## Supplementary Material

### 1. Overview of prior methods for cell-type identification

Various methods have been developed for cell type identification. **SNNCliq** [7] computes a similarity graph among cells, referred to as *shared nearest neighbor (SNN)*. It then uses a graph-based clustering algorithm to identify dense subgraphs. **Seurat** [15] was originally designed for spatial reconstruction of scRNA-Seq data. Since then, it has been extensively updated and used for cell-type identification. In more recent versions (v2.2), Seurat adopted a graph-based approach similar to SNNCliq with extensive modifications that deviate from the original version. **TSCAN** [9] starts by grouping genes with similar expression patterns into “modules” and represents all cells in this reduced space. It then performs principal component analysis (PCA) over the module space to further reduce dimensions. Finally, cells are clustered by fitting a mixture of multivariate normal distributions to the data, with the number of components estimated using the Bayesian Information Criterion (BIC). **SCUBA** [5] first uses k-means with gap statistic to cluster data along an initial binary tree by analyzing bifurcation events for time-course data. Then, it refines the tree using a maximum likelihood scheme. **BackSPIN** [16] is based on the SPIN algorithm, which permutes correlation matrix of cell types to extract its underlying structure. BackSPIN then couples it with a divisive splitting procedure to identify clusters from the ordered similarity matrix. Two methods are specifically designed to identify rare cell types. RaceID [6] uses k-means to first cluster cells, with the number of clusters identified using gap statistic. Then, it identifies rare cell types as outliers that are not explained by an appropriate noise model, accounting for both biological and technical variations. **GiniClust** [38] aims to identify marker genes that are specific to rare cell types using the concept of Gini index. Then, it computes distances between cell types in this reduced subspace and uses DBSCAN clustering algorithm to identify cell types. In addition to these methods, there are approaches that visualize cell types on a continuous spectrum in a given space. Haghverdi *et al.* [39] use diffusion maps to model the continuous spectrum of cells. In another direction, Korem *et al.* [8], adopted a previously developed method, called **Pareto task inference (ParTI)** [17] and applied it to single cell datasets. The latter method itself is based on the original work of Shoval *et al.* [11]. While ParTI uses a similar notion non-convex archetypal analysis as what we do, our method begins with the separable NMF method, the solution of which can be formulated as a convex problem, to pick “ideal” candidate cells as archetypes. Then, it uses the non-convex PCHA procedure to refine these primary archetypes by sparse local averaging to combat noise in the data. Furthermore, our method is founded on a biologically-inspired, kernel-based approach, has a novel method to identify the number of cell types, and last but not the least, a statistical method to construct regulatory circuits that uniquely distinguish each cell type.

## 2. Comparison of Entropy-based marker detection method with Gini index and dispersion

In order to compare the performance of different methods to identify cell type-specific genes, we focused on the **Melanoma** and the **MouseBrain** datasets, for which the original paper provided curated markers for cell types. For each dataset, we ranked genes according to each measure, both before and after adjustment for the effect of universally-expressed genes. Then, for each cell type, we created a true-positive vector based on its curated markers and assessed the over-representation of these markers among top-ranked genes from each method. Finally, we combined each of these over-representation *p*-values using Fisher’s method. Figure 8 illustrates the results for each dataset. As can be seen from the figure, in both datasets, the *dispersion* method is superior before adjustment, whereas Gini index and Entropy-based methods excel after adjustment. As a general trend, we observe that dispersion methods outperform in predicting markers for the most frequent cells in the dataset (T and tumor cells in the **Melanoma** dataset, and S1Pyramidal, CA1Pyramidal, and Oligodendrocyte in the **MouseBrain** datasets), whereas the other two methods significantly outperform dispersion for the rest of cell types, including rare cell types.

**Figure 8:**
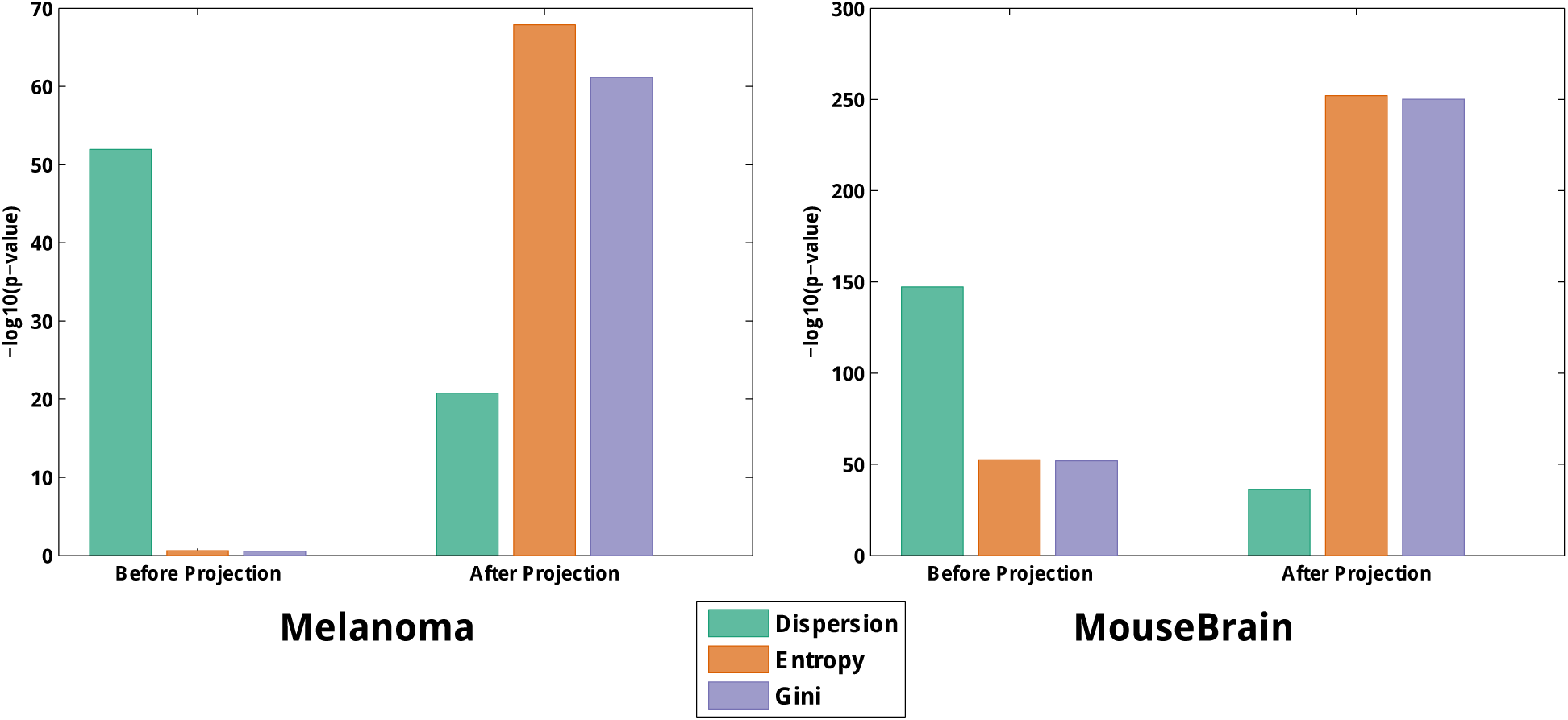
Performance of different marker detection methods before/after correction for the effect of universally expressed genes.

Next, to evaluate the extent of overlap among top-ranked genes, we focused on the top 1,000 genes in each method. Figure 9 shows the Venn diagram for the overlap of datasets. The Gini index and entropy based methods have the highest agreement with each other, while the entropy-based method has a higher overlap with dispersion method than Gini index.

**Figure 9:**
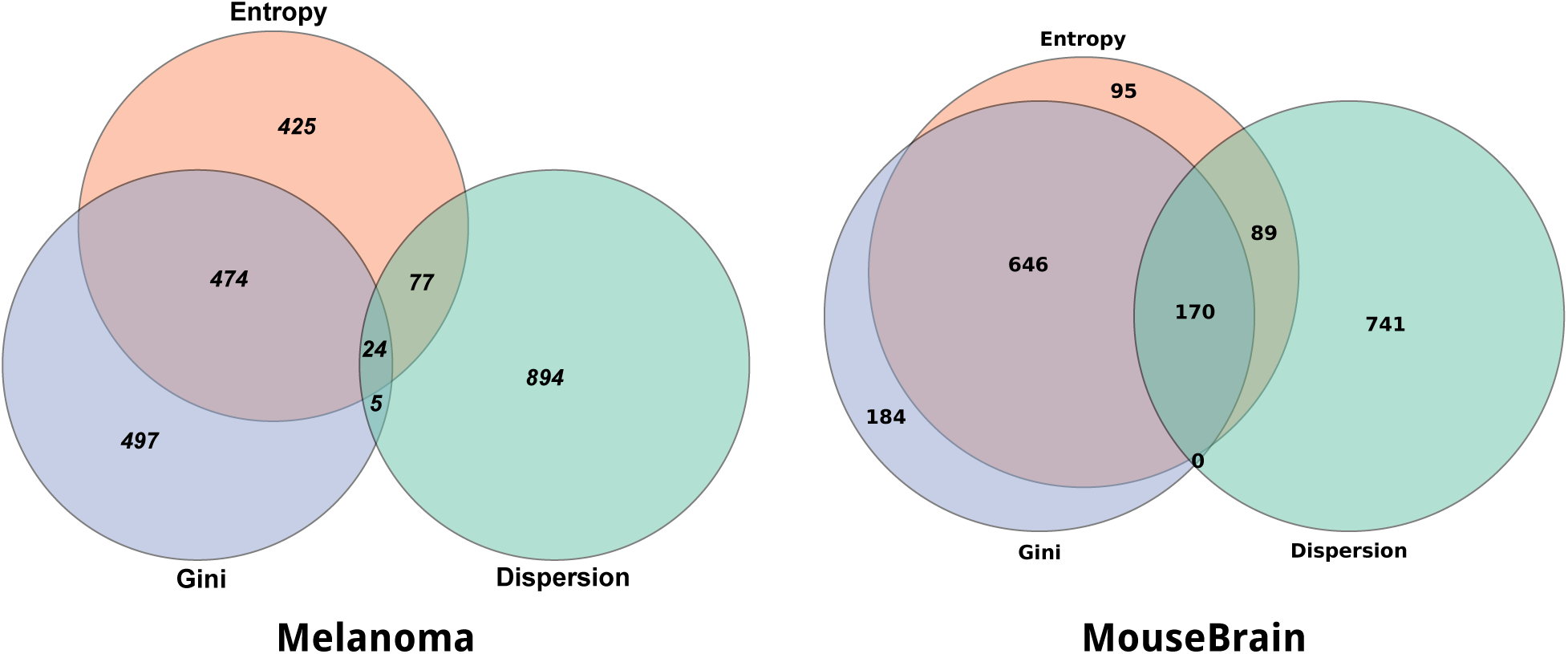
Overlap among top-ranked 1,000 genes predicted using dispersion, Gini index, and entropy-based methods.

## 3. Distribution of clustering measures and significance of differences between different cell similarity metrics

In Figure 3 in the main text, we reported a mean over 100 trials of kernel k-means with the four kernels: ACTION, IsoMap, MDS, and SIMLR. Figure 10 shows the actual distribution of different quality measures for each kernel k-means run. The figure also reports a t-test between the first and second-best method. As in the main figure, ACTION performs equally well or better than other metrics, with the only exception being F-score for the **Brain** dataset.

**Figure 10:**
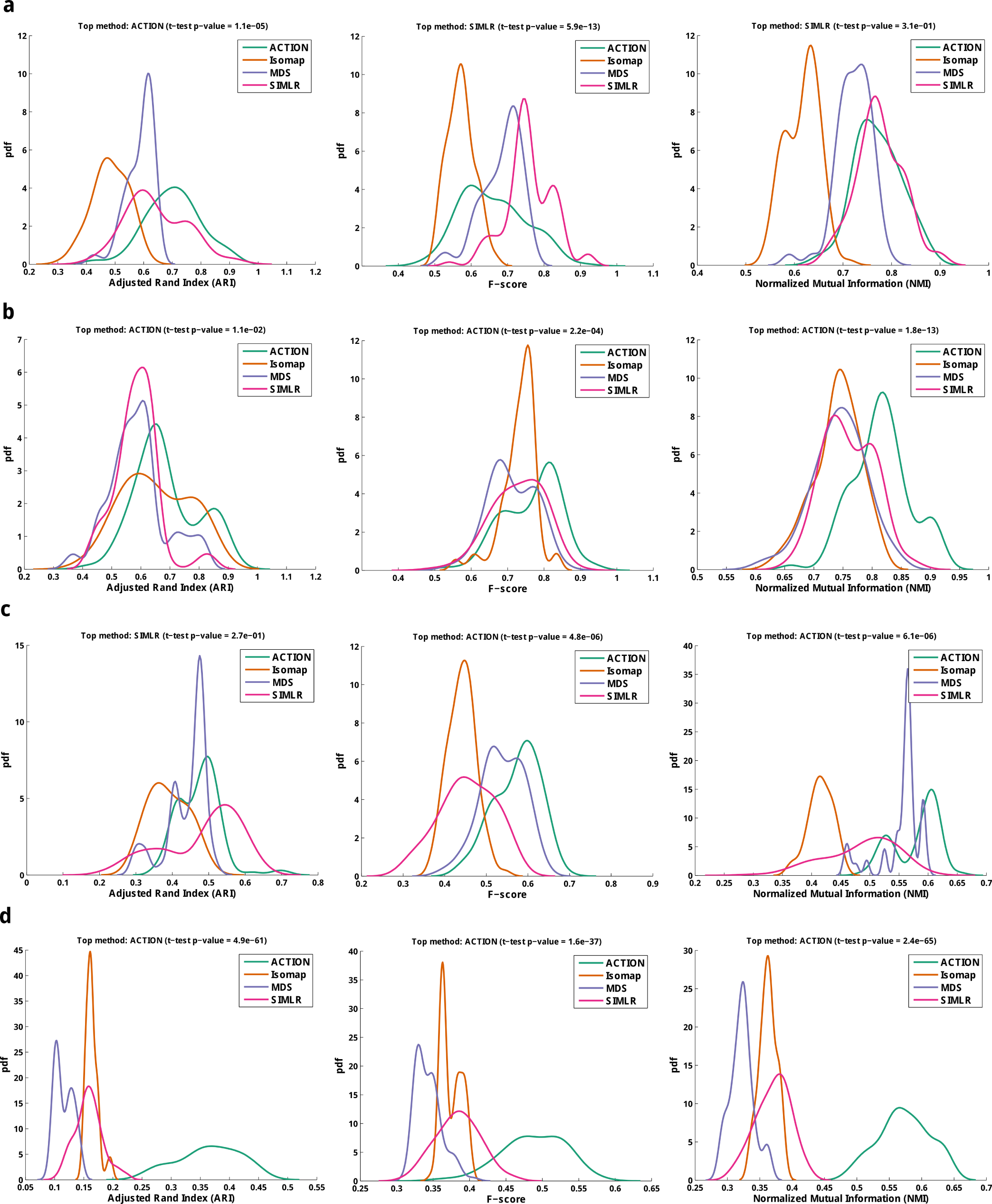
Performance of cell similarity metrics For each extrinsic measure on each dataset, the distribution of values for kernel k-means runs is presented. In each case, the *p*-value of *t*-test between the top-ranked versus runner-up methods has been reported. (**a**) **Brain** dataset, (**b**) **CellLines** dataset, (**c**) **Melanoma** dataset, (**d**) **MouseBrain** dataset.

## 4. Detailed analysis of cell types identified using different similarity metrics – case study in the CellLines dataset

The CellLines data contains measurements from seven distinct cell-lines: A549, GM12878, H1, H1437, HCT116, IMR90, K562. We used this dataset to assess the results from kernel k-means for all of the different metrics. The goal was to use a standard algorithm and compare the results as we vary the type of cell-similarity. Figure 11 shows the subspace of cell line-specific markers, sorted according to the identified cell types in different methods. In all cases, there exists a predicted cell type that mistakenly mixes samples from the *H1437* cell line with one or more other cell lines. In case of *ACTION*, it almost identifies *H1437* perfectly, with marginal contamination from from the *K562* and *IM90* cell lines. This, however, is not surprising since all three of these cell lines are based on lung tissue.

**Figure 11:**
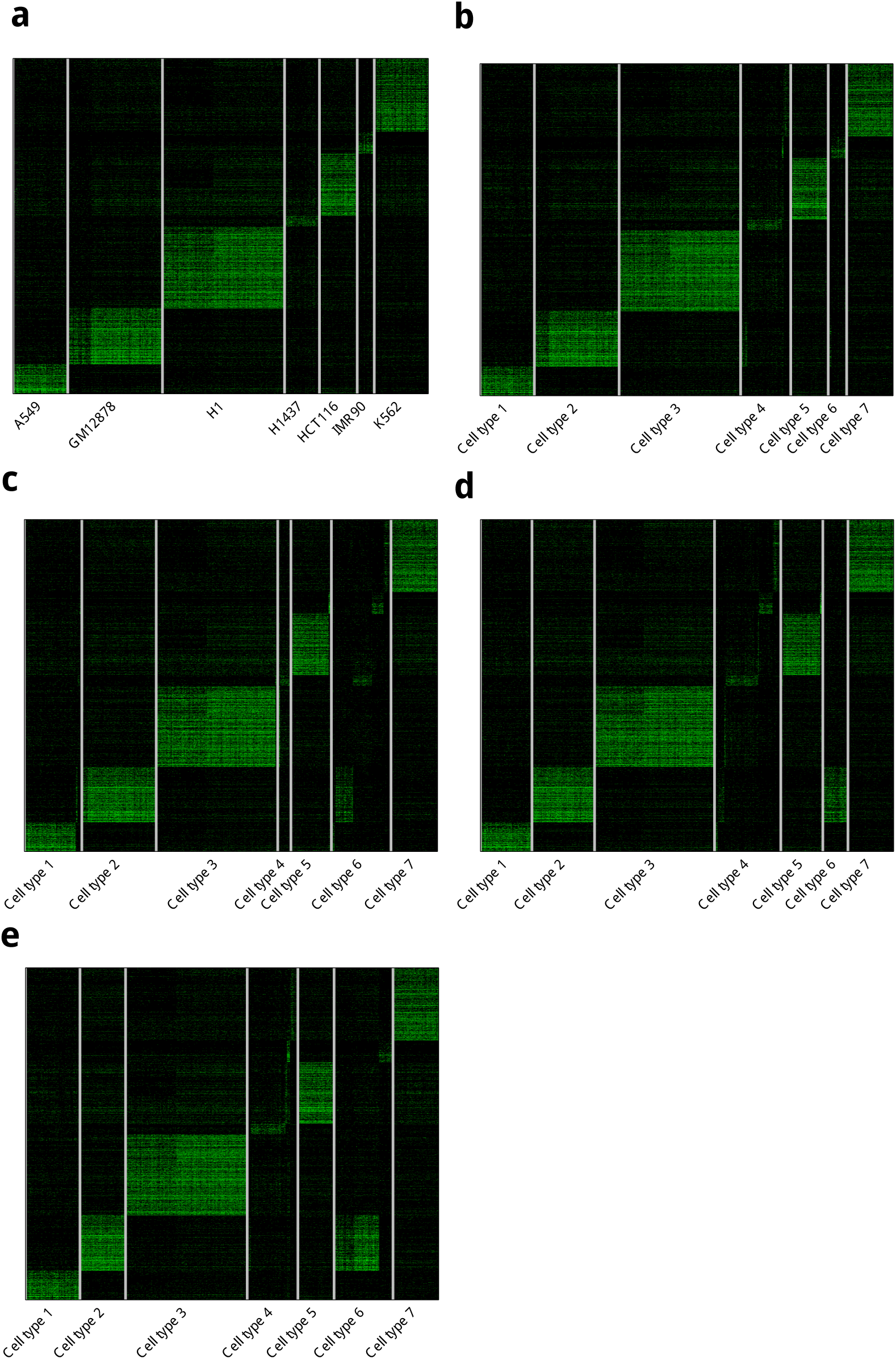
Heatmap of predicted cell types using kernel k-means with different similarity metrics. (**a**) original, (**a**) Original, (**b**) ACTION, (**c**) IsoMap, (**d**) MDS, (**e**) SIMLR

## 5. Detailed analysis of cell types identified using different cell type identification methods – case study in the CellLines dataset

Our next study is similar to the previous one (Supplemental Text 4). In this study, the goal is to compare cell-type identification methods rather than similarity metrics. (Figure 12 shows the marker subspace of identified cell types based on different different cell type identification methods, all of which are non-parametric methods (in the sense that they automatically estimate the number of cell types). Among these methods, *ACTION* that has the highest score and identifies almost all cell types correctly, except that it mixes IMR90 with one of the batches of GM128787. In BackSPIN, all cell lines are either split between two predicted cell types or are mixed with each other. This situation is somewhat better for ParTI, for which the first batch of GM128787 and the H1 cell lines are predicted correctly. However, all other predicted cell types are a mix of different cell lines. SNNCliq splits the cells into too many types. Finally, TSCAN, which performs the second best, mixes parts of K562 and GM128787, and also splits H1 between separate predicted classes. Overall, ACTION shows the highest consistency with the true annotation of cell types, followed by TSCAN.

**Figure 12:**
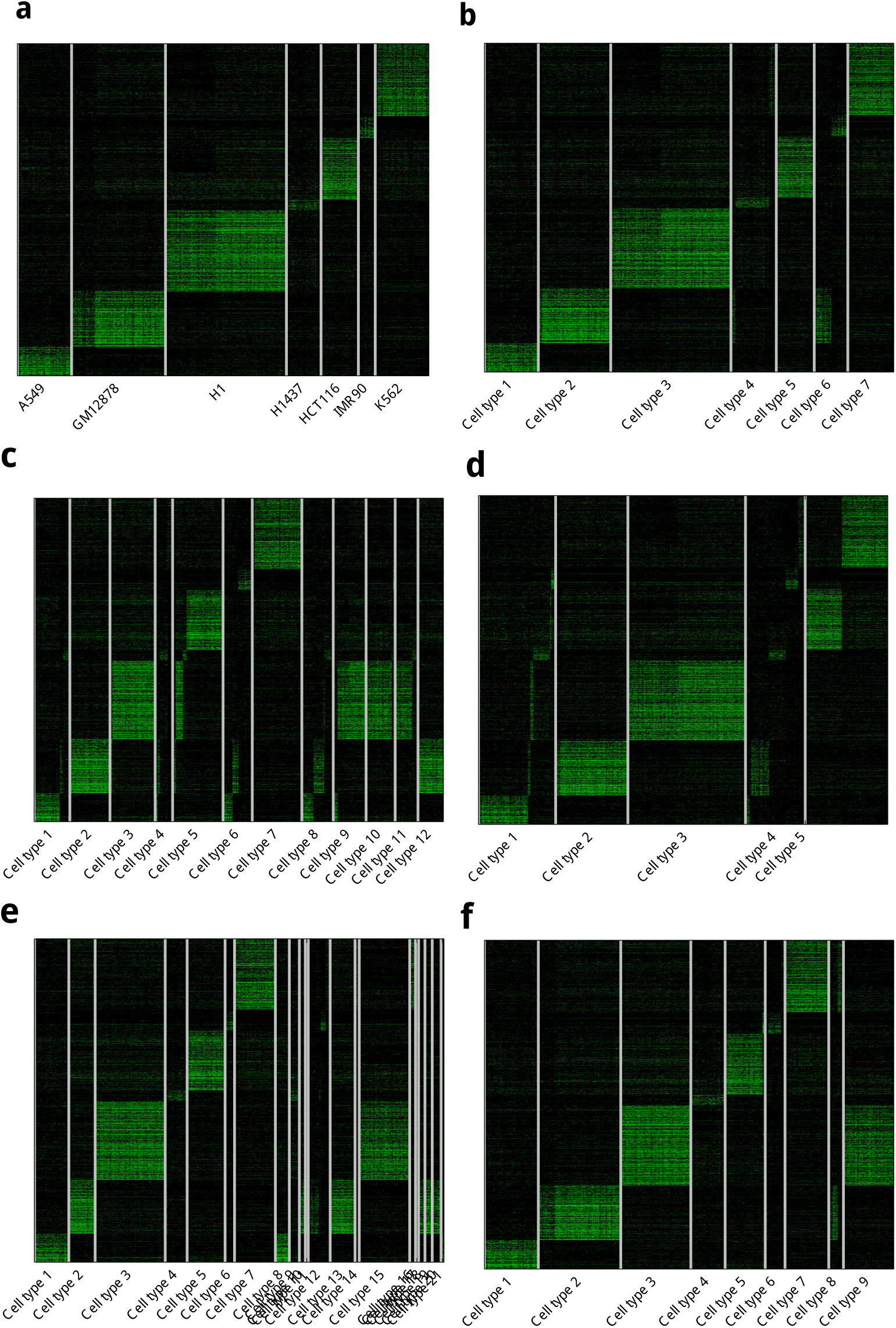
Heatmap of predicted cell types using different cell type identification methods applied to the CellLines dataset. (**a**) original, (**a**) Original, (**b**) ACTION, (**c**) BackSPIN, (**d**) ParTI, (**e**) SNNCliq, (**f**) TSCAN

## 6. Performance of SPA with preconditioner

Let **Y** = **WH**, where matrix **W** is defined as **Y**(:, *𝒮*), with *𝒮* being the selected column subspace of matrix **Y**, and **H** is a non-negative matrix with column-sums equal to one. Moreover, let matrix 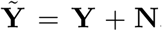, where the noise is bounded: ||**N**(:,*j*)||_2_ ≤ *∊*. Then, the performance of the SPA algorithm has the following upper bound guarantee:

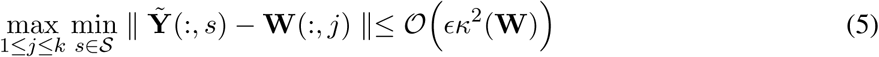

More recently, other techniques have been developed to enhance the robustness of *SPA* to noise [40]. These methods are based on the fact that premultiplying matrix **Y** by a nonsingular matrix **Q** preserves its separability. In this case, the upper bound limit changes to: 𝒪(*∈κ*(**W**)*κ*_3_(**QW**)) Thus, by carefully choosing matrix **Q**, we can enhance the conditioning of the problem. Ideally, if **Q** = **W**^−1^, then *κ*^3^(**QW**) = 1 and we reduced the upper bound from quadratic to linear. While **W**^−1^ is not accessible, we can approximate **W**^−1^ using a *minimum volume ellipsoid* centered around the origin that contains all columns of the original matrix **X**. Formally, this can be solved using the following SDP to identify matrix **A***\**:

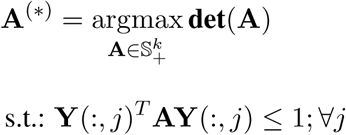

Since **A^T^** is symmetric positive definite, we compute **A^T^** = **Q**^*T*^ **Q** using Cholesky factorization and use it as a preconditioner.

## 7. Pseudo-code for fitting a geometric construct over single cells

See Algorithm 1.

### Algorithm 1 SPA algorithm with prewhitening

**Input: Y** *∈*ℝ^*m×n*^: adjusted expression profile of cells

**Output: A** *∈*ℝ^*m×k*^: primary functions, 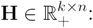 functional identity of cells

1: Solve **minimum volume ellipsoid** problem to identify preconditioner **Q**.

2: **K** = **Y**^*T*^ **Y**, **R** = **QY**, *𝒮* = {}

3: **for** *i* = {1, *…, max_k_*} **do**

4: *α* = argmax_*j*_ ||*r*_j_||_2_{*r*_*j*_ is the *j*th column}

5: *β* = **R**(:, *α*)

6: 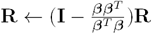 {Orthogonal Projection}

7: *𝒮* ← *𝒮 ∪* {*β*}

8: Construct archetype similarity graph from **G** = **K**(*𝒮, 𝒮*)

9: **if** *subgraph*_*density*_(**G**) is significant **then**

10: **break**

11: **end if**

12: **end for**

13: Initialize **C**_0_ using selected columns in *𝒮*, and run kernel **PCHA** with **K** to estimate matrices **C** and **H**

14: **A** = **YC**

## 8. Computational runtime analysis

In terms of timing, the most time-consuming part of *ACTION* is the preconditioning using minimum volume ellipsoid method, which depends on the solver being used. Using CVX with Mosek solver, timings are as reported in Figure 13. For larger datasets, it can be seen that *ACTION* scales more gracefully compared to other methods.

**Figure 13:**
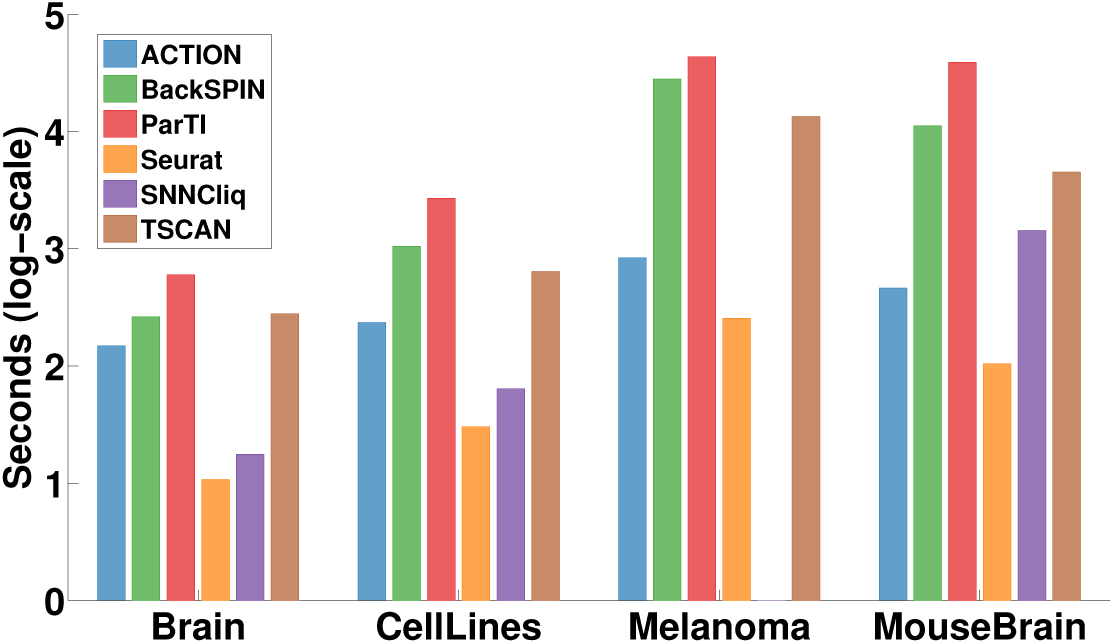
Time-wise, smaller values are the better. It can be seen that *ACTION* scales better as the datasets grow.

## 9. Robustness of *ACTION* method in presence of noise and outliers

To further evaluate the effect of preconditioning convex Non-negative Matrix Factorization (NMF), as well as relaxing it with Principal Convex Hull Analysis (PCHA), we performed a simulation to assess the impact of outliers on these methods, as well as to find the critical point at which an outlier becomes a rare cell type. To this end, we again focus on the **CellLines** dataset. In this case, *H1* cell line (embryonic stem cell) is the farthest from the rest of cell lines. We set up an experiment in which we held out *H1* and gradually introduced different percentages of *H1* cells, varying from one to ten percent. For each case, we tried 10 individual replicas. Figure 14a-c presents the performance of each method in identifying cell types, measured with respect to known cell types. In each case, we observe that preconditioning (Pre-SPA) significantly enhances the quality of results compared to Successive Projection Algorithm (SPA) alone. However, this makes the results unstable (has a high variance). Applying PCHA on top (PreSPA+PCHA) smooths out these variations. In order to assess the performance of these methods in identifying rare cell types, we used bipartite matching in each case to find the closest predicted cell type to *H1* and then used hypergeometric *p*-value to assess the overlap of these two sets. These results, presented in Figure 14d, show that both PreSPA, and PreSPA+PCHA are sensitive enough to identify rare cell types. However, PreSPA is sensitive to low percentages of introduced *H1*, whereas PreSPA-PCHA considers percentages less than 2% to be noise/outlier and after that starts to identify it as a rare cell type.

**Figure 14:**
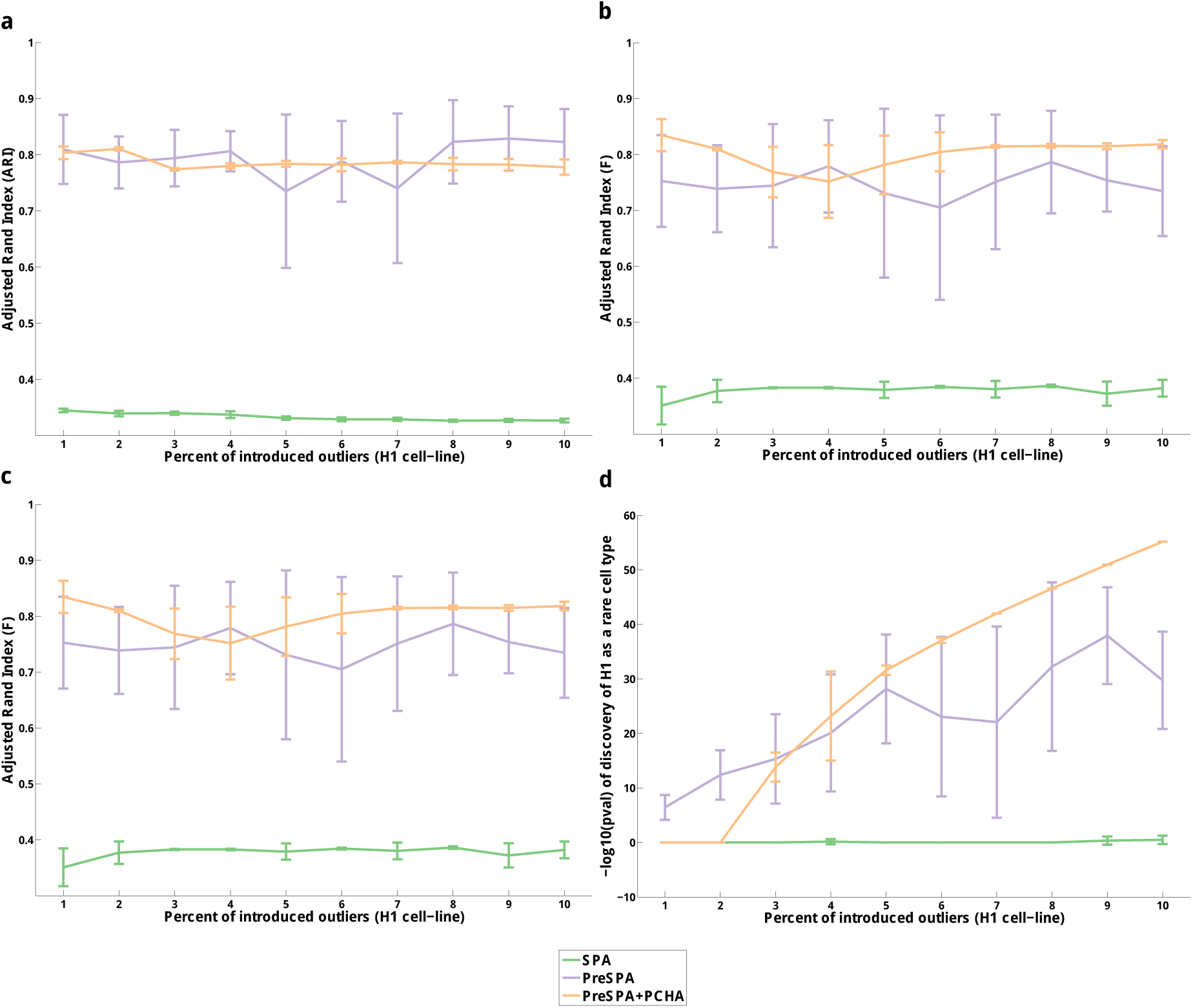
Robustness of *ACTION* method in presence of noise and outliers. **(a)-(c)** Different measures of cell type identification quality as a function of introduced noise, **(d)** Analysis of the critical point of transitioning from noise to rare cell type.

## 10. Visualizing the functional space of cells

Unlike the conventional application of t-distributed stochastic neighbor embedding (tSNE), which is used to project the transcriptional profile of cells into a lower dimensional space, we propose a framework that captures the distribution of cells around archetypes. To this end, we focus on the functional identity of cells with respect to our archetypes, which is computationally represented by the matrix **H**. Each column in this matrix is a stochastic vector (sums to one) that represents the extent to which a cell is close to a given archetype. To visualize this continuous functional space, we first initialize the solutions using the Fiedler embedding (as opposed to tSNE that uses random initialization). Then, we use tSNE to update the initial coordinates. The following pseudo-code illustrates the proposed projection.

1. Take **H** from the PCHA as input.
2. Set 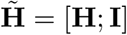
3. Let 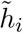 be the *i*th column of 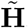, and compute entries of the matrix 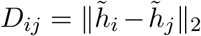 (that is, Euclidean distance between vectors 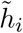 and 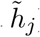).
4. Convert Distances to Similarity following Network Similarity Fusion [41] affinity matrix construction

a. Let 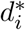 be the average distance from *i*^*th*^ cell to its top *k* = **round**(*n/*10) closest neighbors, with *n* being the total number of cells. (If you sort columns of the matrix **D**, this is just the top *k* entries.)
b. Set 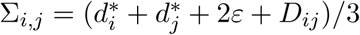, where *ε* is 2^−52^.
c. Set 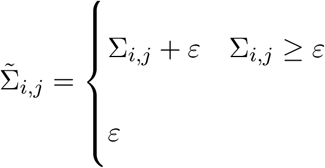
d. Set *W*_*i,j*_ to be the probability that a normally distributed random variable with mean 0 and standard deviation 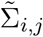 has value *D*_*i,j*_.
5. Set **G** = (**W** + **W**^*T*^) /2 be the weighted graph between cells.
6. Set **L** = diag(**G** *·* ones(*n*, 1)) – **G** (that is, **L** is the combinatorial Laplacian of **G**).
7. Compute the three smallest eigenvalues and eigenvectors of **L**, (*υ*_1_, *λ*_1_), (*υ*_2_, *λ*_2_), (*υ*_3_, *λ*_3_). Note that *λ*_1_ is zero because of the Laplacian structure.
8. Set *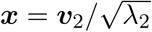*
9. Set *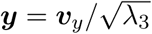*
10. Run t-SNE to update *x, y* coordinates.
11. Final map represents the distribution of cells around each archetype.

## 11. List of 20 top-ranked genes for each archetype in the Melanoma dataset

See Table 1

**Table 1:** Table of the top 20 residual genes after orthogonalization. Each archetype is also annotated with its enriched cell type. Bolded genes are the genes that coincide with known markers provided by the original paper. The last row is the *p*-value of enrichment of markers among all genes sorted after orthogonalizing each archetype.

| TF | *Subclass B* | *Subclass C* |
| --- | --- | --- |
| MITF | 0.00E+00 | 2.14E-04 |
| E2F1 | 5.18E-06 | 7.62E-01 |
| LEF1 | 3.19E-04 | 3.83E-01 |
| MYC | 8.39E-04 | 3.03E-01 |
| TFEB | 2.30E-04 | 2.63E-02 |
| E2F4 | 1.32E-02 | 5.11E-01 |
| TCF7 | 3.32E-02 | 1.00E+00 |
| CNBP | 4.05E-02 | 1.00E+00 |
| FUBP1 | 4.05E-02 | 1.00E+00 |
| RBL1 | 4.05E-02 | 1.00E+00 |
| SRSF1 | 4.05E-02 | 1.00E+00 |
| TBL1X | 4.05E-02 | 1.00E+00 |
| SNIP1 | 1.30E-02 | 2.90E-01 |
| MTA1 | 4.05E-02 | 6.36E-01 |
| HOXA1 | 1.30E-02 | 1.96E-01 |
| OTX2 | 3.32E-02 | 1.71E-01 |
| ONECUT2 | 4.05E-02 | 1.31E-01 |
| SOX9 | 2.84E-03 | 4.57E-03 |
| NFIB | 3.60E-01 | 3.14E-02 |
| TBP | 4.62E-01 | 2.52E-02 |
| USF1 | 4.62E-01 | 2.49E-02 |
| KLF5 | 7.85E-01 | 3.97E-02 |
| POU4F2 | 3.60E-01 | 1.73E-02 |
| ATF3 | 1.00E+00 | 4.54E-02 |
| PAX3 | 3.59E-01 | 1.62E-02 |
| ESR1 | 1.07E-01 | 4.57E-03 |
| MYF6 | 1.00E+00 | 4.02E-02 |
| HDAC7 | 1.00E+00 | 3.97E-02 |
| MAML1 | 1.00E+00 | 3.97E-02 |
| NKX2-3 | 1.00E+00 | 3.97E-02 |
| PPARG | 1.00E+00 | 3.87E-02 |
| LCOR | 6.33E-01 | 2.45E-02 |
| SMAD4 | 7.85E-01 | 2.63E-02 |
| GTF2I | 1.00E+00 | 3.14E-02 |
| TAF1 | 1.00E+00 | 3.00E-02 |
| SOX10 | 1.03E-02 | 3.00E-04 |
| ETS2 | 9.57E-01 | 2.63E-02 |
| ETS1 | 1.00E+00 | 2.53E-02 |
| MYF5 | 1.00E+00 | 2.49E-02 |
| HOXA7 | 8.15E-01 | 1.50E-02 |
| EGR2 | 8.24E-01 | 1.39E-02 |
| STAT3 | 1.00E+00 | 1.47E-02 |
| JUND | 9.48E-01 | 1.39E-02 |
| KLF4 | 1.00E+00 | 1.24E-02 |
| TWIST2 | 6.52E-01 | 7.91E-03 |
| KLF6 | 8.10E-01 | 8.97E-03 |
| TP53 | 3.11E-01 | 3.06E-03 |
| TFCP2 | 8.15E-01 | 7.30E-03 |
| ESR2 | 1.00E+00 | 7.30E-03 |
| PARP1 | 1.00E+00 | 7.30E-03 |
| ETV4 | 1.00E+00 | 4.57E-03 |
| PPARD | 1.00E+00 | 4.57E-03 |
| CTNNB1 | 3.50E-01 | 1.58E-03 |
| AR | 1.00E+00 | 1.58E-03 |
| BRCA1 | 1.00E+00 | 2.14E-04 |

| 1-T | 2-B | 3-T/Unresolved | 4-Tumor | 5-Tumor | 6-Tumor | 7-Macro | 8-Endo/CAF |
| --- | --- | --- | --- | --- | --- | --- | --- |
| **NKG7** | **MS4A1** | UGDH-AS1 | DCT | APOC2 | SAA1 | **TYROBP** | IGFBP7 |
| **CD8A** | **CD79A** | ROCK1P1 | **PMEL** | APOD | TF | **FCER1G** | EFEMP1 |
| **CST7** | HLA-DRA | TMEM212 | LHFPL3-AS1 | **SERPINA3** | SFRP1 | **CD14** | CCL21 |
| GZMA | **BANK1** | HERC2P4 | CTSK | **MIA** | MAGEA4 | **IFI30** | BGN |
| **CD3D** | **CD79B** | ASTN2 | TYRP1 | A2M | RGS5 | **C1QC** | **THY1** |
| CCL4 | IGLL5 | SHISA9 | TUBB4A | **SERPINE2** | MAGEC2 | **C1QA** | TFPI |
| **IL32** | **IRF8** | ORC4 | SCD | **TYR** | PDK4 | **AIF1** | CLDN5 |
| **GZMK** | **CD37** | SPC25 | **GPM6B** | APOE | C2orf82 | **C1QB** | PDLIM1 |
| **PRF1** | **CD19** | LOC643406 | **RAB38** | IFI27 | ALDH1A3 | **S100A9** | **COL1A1** |
| **CD2** | CD74 | LOC646214 | **PIR** | TRIML2 | CAMP | **FCGR3A** | **C1R** |
| **KLRK1** | CXCR4 | ODF2L | KIT | MFGE8 | **SERPINA3** | **CSF1R** | **RARRES2** |
| ITM2A | SELL | ABCC9 | CA14 | NSG1 | C1QTNF3 | **MS4A6A** | CYR61 |
| RGS1 | **VPREB3** | L2HGDH | BCAN | RDH5 | ANGPTL4 | **IGSF6** | **DCN** |
| PDCD1 | HLA-DPA1 | LYZ | SNAI2 | **SLC26A2** | ERRFI1 | **HCK** | **C1S** |
| CD27 | TCL1A | LOC286437 | GSTO1 | **CAPN3** | COL1A2 | **PILRA** | NNMT |
| **TIGIT** | HLA-DQB1 | MAB21L3 | **MLANA** | MT2A | MRPL36 | **VSIG4** | CXCR7 |
| **LCK** | **BCL11A** | KCNQ1OT1 | **SLC45A2** | SPP1 | FN1 | IL1B | IGFBP4 |
| CTSW | LTB | ARHGEF26-AS1 | **GPR143** | TM4SF1 | HAPLN1 | TMEM176B | ECSCR |
| IL2RG | NAPSB | FBLIM1 | TRPM1 | **PMEL** | SAA2 | **CD163** | GNG11 |
| **SIRPG** | **CD22** | GLIPR1L2 | **CDK2** | **CDH19** | MGP | FCN1 | CLU |
| $7.7 \times 10^{-67}$ | $3.5 \times 10^{-59}$ | $1.6 \times 10^{-3}$ | $2.5 \times 10^{-22}$ | $4.8 \times 10^{-53}$ | $5.0 \times 10^{-05}$ | $2.4 \times 10^{-176}$ | $5.9 \times 10^{-98}$ |

## 12. List of functionally active transcription factors

The following table lists all transcription factors (TFs) that are either significant in *subclass B, archetype 5, subclass C, archetype 4*, or both. Second and third columns of the table are the *p*-values of the functional activity of TFs. These factors are sorted according to the relative importance in these subclasses *B* and *C*. Green rows are the ones that are significant in both. *TP53* and *MYC*, marked in red, are used in conjunction with *MITF* to distinguish these two classes.

## 13. Regulated downstream targets of MITF factor in Subclasses *B* and *C*

The following table lists the full set of significant downstream targets of MITF in both subclasses *B & C*. Genes *GPNMB, MLANA, PMEL* and *TYR* are shared between two subclasses, whereas the rest of targets are unique to one of them. For genes that have significant effect on the survival rate, their Cox coefficient is presented in the table. A positive Cox coefficient indicates that high expression of the given genes is associated with poor survival.

| Target | Subclass B | Subclass C | Cox Coefficient |
| --- | --- | --- | --- |
| ACP5 | ✓ |  | - |
| CDK2 | ✓ |  | 0.218 |
| CTSK | ✓ |  | - |
| DCT | ✓ |  | - |
| KIT | ✓ |  | 0.3214 |
| OCA2 | ✓ |  | 0.3038 |
| TRPM1 | ✓ |  | 0.188 |
| TYRP1 | ✓ |  | 0.2422 |
| GPNMB | ✓ | ✓ | - |
| MLANA | ✓ | ✓ | - |
| PMEL | ✓ | ✓ | 0.2765 |
| TYR | ✓ | ✓ | - |
| BEST1 |  | ✓ | - |
| BIRC7 |  | ✓ | - |
| FOS |  | ✓ | - |
| MET |  | ✓ | - |

## 14. Regulatory networks

We provided transcriptional regulatory networks (TRNs) for all eight archetypes in the Melanoma dataset. There are two files per network, one edge list file and one node annotation file, both in plain text (txt) format. The former contains edges in form of “TF name tab TG name,” whereas in the second file there is a row for each node (either TF or TG) that provides additional information about it. This information includes: (i) type (TF or TG), (ii) -log10 of the functional activity p-value, if node is a TF, or zero otherwise, (iii) residual expression of genes (either TF or TG) after orthogonalization of archetype, (iv) heuristic importance of node (for visualization purposes only), and (v) cox survival coefficient.

